# Effective online Bayesian phylogenetics via sequential Monte Carlo with guided proposals

**DOI:** 10.1101/145219

**Authors:** Mathieu Fourment, Brian C. Claywell, Vu Dinh, Connor McCoy, Frederick A. Matsen, Aaron E. Darling

## Abstract

Modern infectious disease outbreak surveillance produces continuous streams of sequence data which require phylogenetic analysis as data arrives. Current software packages for Bayesian phylogenetic inference are unable to quickly incorporate new sequences as they become available, making them less useful for dynamically unfolding evolutionary stories. This limitation can be addressed by applying a class of Bayesian statistical inference algorithms called sequential Monte Carlo (SMC) to conduct *online inference*, wherein new data can be continuously incorporated to update the estimate of the posterior probability distribution. In this paper we describe and evaluate several different online phylogenetic sequential Monte Carlo (OPSMC) algorithms. We show that proposing new phylogenies with a density similar to the Bayesian prior suffers from poor performance, and we develop ‘guided’ proposals that better match the proposal density to the posterior. Furthermore, we show that the simplest guided proposals can exhibit pathological behavior in some situations, leading to poor results, and that the situation can be resolved by heating the proposal density. The results demonstrate that relative to the widely-used MCMC-based algorithm implemented in MrBayes, the total time required to compute a series of phylogenetic posteriors as sequences arrive can be significantly reduced by the use of OPSMC, without incurring a significant loss in accuracy.

## Introduction

Phylogenetic techniques are quickly becoming an essential tool in the investigation and surveillance of infectious disease outbreaks (Rusu et al. 2015; Gardy et al. 2015; Neher and Bedford 2015). Meanwhile, advances in DNA sequencing technology have made the generation of complete genome data for isolates of bacteria and viruses a routine practice in public health laboratories. These genome data are collected into public databases such as the FDA GenomeTrakr (FDA 2016), which in 2016 accumulated new data at an average rate of over 1000 pathogen genomes per week. Sequencing technology itself continues to evolve, with new devices based on nanopore detection capable of generating a continuous stream of sequence data, supporting interactive real-time analysis (Loose et al. 2016).

Ideally these new data streams would be matched with appropriate sequence analysis tools, including Bayesian phylogenetic inference. Bayesian inference has particular value in epidemiological investigations due to its ability to operate on models with a wide range of unknown parameters, including divergence times, lineage-specific mutation rates, population demographics, and geography (Kühnert et al. 2014; Lemey et al. 2009). However, all current methods for Bayesian inference treat the data set as a static entity that has been observed in its entirety at the time that computation of the posterior probability distribution begins. Updating a dataset with new sequences, as might be required when a new case of an infection is presented and sequenced, necessitates that the entire analysis be restarted.

Although Izquierdo-Carrasco et al. (2014) have proposed a maximum likelihood approach to update a phylogenetic tree with new sequences, no such tool exists for Bayesian phylogenetic inference. Each run using popular Bayesian phylogenetic inference tools like MrBayes (Ronquist et al. 2012) or BEAST (Bouckaert et al. 2014) can take days or weeks of CPU time to approximate a posterior distribution for realistic models and datasets. The inability to quickly incorporate new data into an existing analysis is a major impediment to the use of Bayesian phylogenetics as a decision support tool for infectious disease management and surveillance, where interventions are most likely to be effective if made within hours or days.

Recently, Dinh et al. (2016) described a theoretical framework for updating a phylogenetic posterior approximation, called Online Phylogenetic Sequential Monte Carlo (OPSMC). An overview of OPSMC is given in Figure 1. At each iteration, a transition kernel is used to update the posterior on *n –* 1 sequences to approximate the posterior on *n* sequences. Optionally, one or more Metropolis-Hastings steps (not shown in the figure) can be applied to increase the effective sample size. Although these authors showed the OPSMC framework has attractive theoretical properties, it was not clear from that work whether it can be translated into an effective sampler. In addition, more research is needed on the design of effective transition kernels for OPSMC, a subject of some debate in related literature (Teh et al. 2008; Bouchard-Côté et al. 2012).

**Figure 1.**
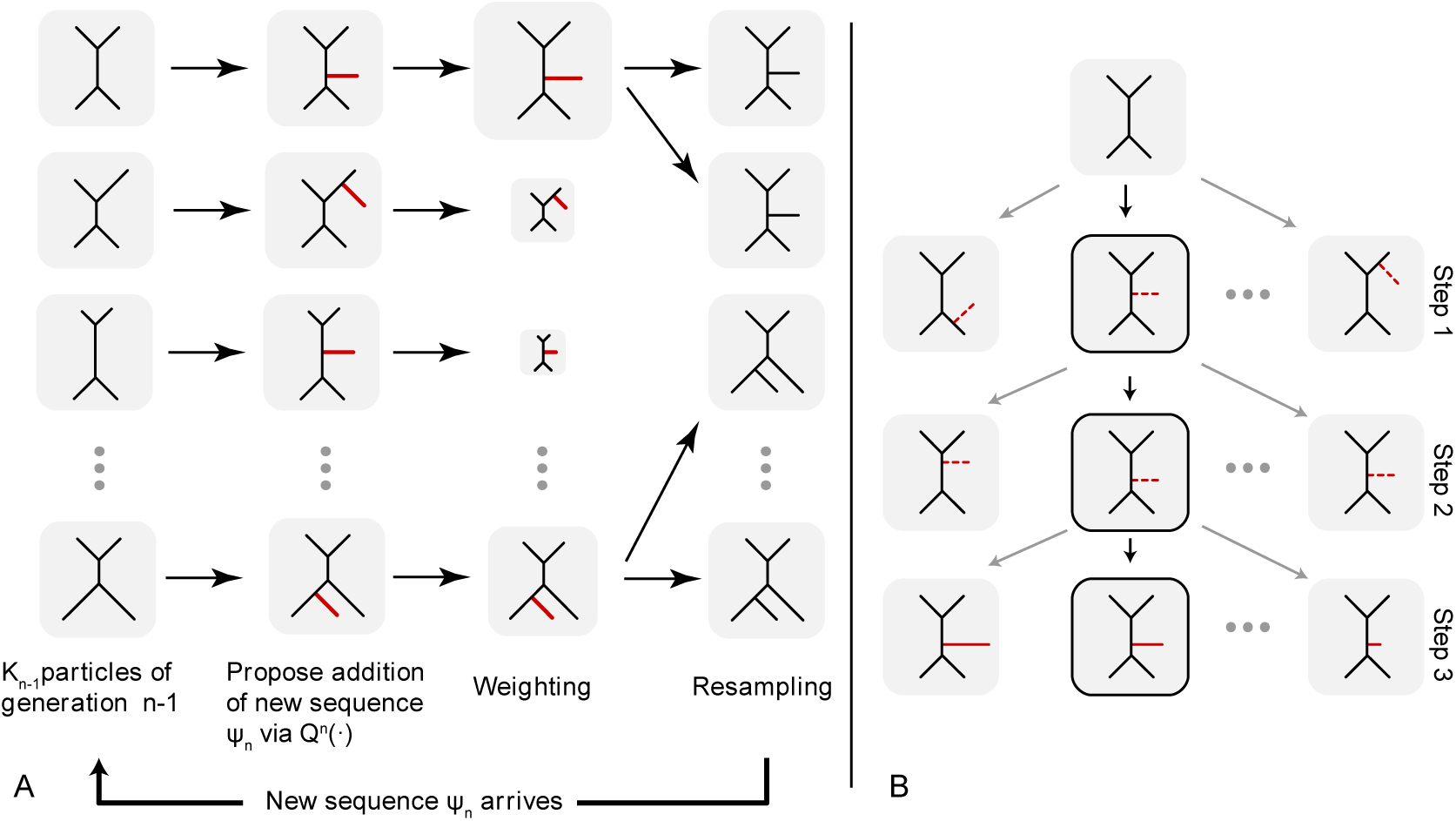
An overview of the Online Phylogenetic Sequential Monte Carlo algorithm as implemented in `sts`. Panel A shows a population of particles going through an SMC iteration. For each particle, a new sequence (taxon) *ψ*_*n*_ is attached to the tree using proposal *Q^n^*(•), its weight is computed and particles are resampled using the weights. Panel B depicts the three-step proposal applied by `sts` to a single particle.

In this work we implement OPSMC with a variety of transition kernels, and compare their ability to efficiently update phylogenetic posteriors with new data. In particular we compare the efficiency of naïve proposals to guided proposals, showing that the extra effort required to compute a guided proposal leads to a significant overall improvement in sampler efficiency Finally, we discuss prospects for the incorporation of OPSMC into widely used algorithms and software packages for Bayesian phylogenetic inference. For this paper, we restrict ourselves to “pure” SMC without Metropolis-Hastings steps. Our implementation is available at https://github.com/OnlinePhylo/sts/.

Our results build upon several key pieces of previous work building trees using SMC via subtree merging. Teh et al. (2008) were the first to describe the use of Sequential Monte Carlo for Bayesian inference of tree-structured models. Bouchard-Côté et al. (2012) adapted that work to infer rooted, ultrametric phylogenetic trees. Wang et al. (2015) showed that SMC could also be applied to unrooted phylogenetic trees and provided an implementation of the algorithm and performance comparison with the widely used MrBayes software. Because each of these methods proceeds by joining the roots of subtrees merged by previous steps, they can only add additional sequences at the root of the tree. Thus each of those previous contributions is only appropriate for the case where the data set is static and completely known when inference begins.

In contrast, our work relaxes those assumptions to evaluate algorithms for online inference. Dinh et al. (2016) was the first to describe a theoretical framework to extend phylogenetic SMC approaches to online inference. In parallel work, Everitt et al. (2016) have also described an online phylogenetic inference step as part of a larger framework of SMC methods for spaces of varying dimension (see also Everitt et al. 2017). We will compare our work and that of these authors in the Discussion.

## Material and Methods

### Definitions

In Bayesian inference we are interested in estimating the posterior distribution of a model conditioned on data. In phylogenetics, the data are a set of nucleotide or amino acid sequences *ψ* = (*ψ*_1_, *ψ*_2_,…, *ψ*_*N*_) collected from *N* taxa, wherein the homologous nucleotides among sequences have been identified and grouped as sites (columns) in a sequence alignment (Felsenstein 2004). We assume that alignment sites are IID and that mutation events along each branch of a phylogenetic tree *τ* occur independently accordingly to a continuous time Markov chain. In this paper we use the Jukes-Cantor (Jukes et al. 1969) substitution model with equal base frequencies and equal transition rates.

The posterior probability of a phylogenetic tree with topology *τ* and branch lengths l = (*l*_1_, …, *l*_2*N*−3_) conditioned on *ψ* follows from Bayes Theorem:

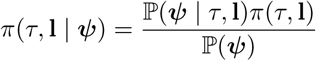

where ℙ(*ψ* | *τ*, l) is the phylogenetic likelihood calculated using the standard Felsenstein pruning algorithm (Felsenstein 2004) and *π*(*τ*, l) is the prior on the topology and branch lengths of the phylogenetic tree. For unrooted trees, branch length priors are usually assumed to be IID with a simple distribution such as truncated uniform or exponential. A prior can also be specified on the unrooted topologies, a common choice being the uniform distribution. The marginal likelihood of the model 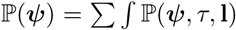 is analytically intractable. Therefore the joint posterior distribution is usually approximated using Monte Carlo methods.

### Sequential Monte Carlo

Sequential Monte Carlo (SMC) algorithms are a class of sampling methods that have been extensively investigated in the context of sequential Bayesian inference. We consider that data arrive sequentially *ψ*_1_, …, *ψ*_*N*_ and we wish to update the approximation of a probability distribution.

The idea is to track, at each iteration n, a collection of *K*_*n*_ particles 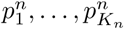 associated with positive weights 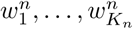 whose empirical distribution converges to the target distribution *π*_*n*_.

Given a collection of weighted particles from the previous iteration *n –* 1 the algorithm applies the three steps: resampling, mutation, and reweighting.

The resampling step prunes particles associated with low weights. This step is optional and is typically triggered when the effective sample size (Beskos et al. 2014) of the particle collection drops below a predetermined threshold. The effective sample size (ESS) for a collection of particles with weight 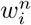 for the *i*th normalized particle at iteration *n* is

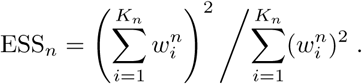

Resampling obtains *K*_*n*_ particles, 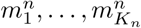, via a multinomial distribution on the particles parameterized with the weights 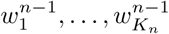. Alternatively, more sophisticated resampling methods such as stratified resampling (Kitagawa 1996) and residual resampling can be used in order to reduce the variance of the new particle population (Doucet et al. 2001; Del Moral et al. 2012).

The mutation step draws *K*_*n*_ new particles from a proposal distribution 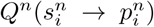 for *i* = 1, …, *K_n_.*

The unnormalized weight 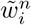 of each particle 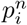 is updated:

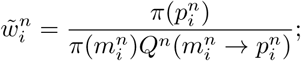

after renormalizing these weights we obtain an empirical approximation of *π*_*n*_:

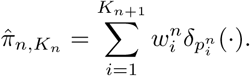

The SMC sampler initializes each particle with equal weights 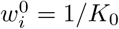 for all i = 1, …, *K*_0_.

### Online phylogenetic sequential Monte Carlo

Given an initial set of phylogenetic trees that represent a sample from the posterior distribution, we set out to update the posterior approximation represented by these samples with new sequences using an Online Phylogenetic SMC (OPSMC).

While the state space of standard SMCs is of fixed dimension, the model complexity and dimension in the OPSMC setting increases as the number of taxa increases. Indeed, the number of tree topologies increases super-exponentially with the number of taxa for both rooted and unrooted trees (St. John 2016). In addition to the discrete component of the tree space, the addition of each taxon requires additional continuous parameters. For rooted trees with a molecular clock, each additional taxon introduces two new parameters (the coalescence time and the identity of the branch where the new lineage attaches) whereas three parameters are introduced in the non-clock case: the attachment branch, the attachment position on the attachment branch, and the length of the pendant branch leading to the new taxon. Unless stated otherwise, in the rest of the paper trees are assumed to be unrooted with no clock. Nevertheless, it will be convenient for the purposes of description that the trees have been given an arbitrary root.

The OPSMC algorithm assumes that sequences arrive sequentially one by one: even if several new sequences have become available, every particle will incorporate the same sequence at a given iteration. This simplification circumvents the over-counting problem highlighted in (Bouchard-Côté et al. 2012; Wang et al. 2015) who showed that uniform tree proposals were biased toward balanced tree topologies. It should be clear that, unlike previously described phylogenetic SMCs, the OPSMC method does not require specifying an extension over a forest of trees since each particle represents a single tree.

Each of the proposals follows a three step process:

1. Choose an attachment branch *e* from 2 *n –* 3 branches.
2. Choose location *x* along *e* to attach the new taxon. We refer to *x* as the *distal length:* distance from the attachment location to the end of the edge that is farthest away from the root of the tree. In these proposals the length of the attachment branch does not change.
3. Propose a new pendant branch length *y* leading to the new taxon.

One can mix and match choices for each of these steps from the following strategies.

We encode each step with a single letter to distinguish among the possible methods used at that step (Table 1). We will use the resulting three-letter code to describe a complete proposal strategy: for example, the LAF proposal uses the L method for the first step, A for the second step, and F for the third step. Some of the methods are decorated with a tilde in order to distinguish heated from unheated proposals (the exact meaning of a heated proposal will be clarified below).

**Table 1.**
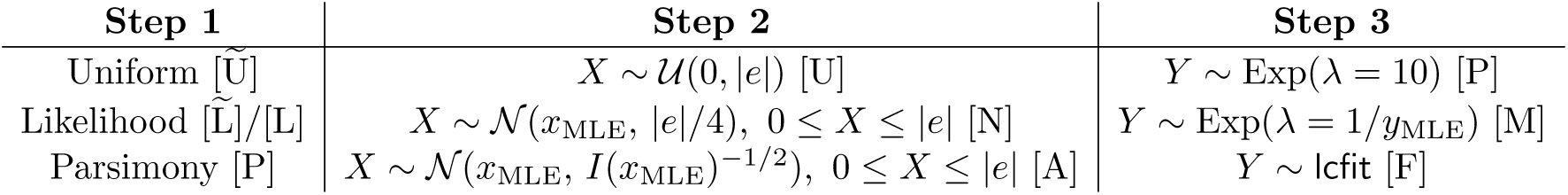
A summary of the proposal distributions for the three steps: first, sampling an attachment edge, second, sampling a position at which to attach the new pendant branch, and third, sampling a pendant branch length. The one-letter code of each sampling strategy is between square brackets. In Step 1, the one-letter code is decorated with a tilde for non-heated proposal.

In the remainder of this section we describe different proposals for each of the three steps and for each step two broad classes of methods are described. The simplest methods are called unguided proposals. Although unguided proposals are fast and simple to implement they tend to generate a very large number of particles with low likelihoods. This causes much CPU time to be expended on calculating likelihoods and SMC weights for particles that ultimately drop out of the posterior approximation during the weighted resampling step. Guided proposals refer to more complex methods that use the data to get more accurate proposals.

#### Step 1: attachment branch choice

##### Uniform 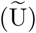 proposals

The unguided uniform proposals are the simplest proposal scheme investigated in this paper and they bear similarities to the “PriorPrior” proposal described in (Teh et al. 2009). In our implementation of uniform proposals the attachment branch is chosen uniformly among all branches. Alternatively, the branch can be selected with weight proportional to its length.

##### Likelihood 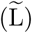 proposals

For each branch in the current tree, an attachment weight for the new taxon is calculated as described below. The attachment branch is then drawn randomly according to a multinomial distribution parameterized with the attachment branch weights.

Calculating the maximum likelihood attachment configuration for each branch is computationally expensive so we instead resort to a heuristic approach inspired by a similar strategy used in pplacer (Matsen et al. 2010). The new taxon is attached in the middle of each branch and likelihoods are calculated using a set of predetermined branch lengths. By default OPSMC calculates the tree likelihood with pendant branch length equal to 0 and separately with the pendant branch length set to the median branch length from the first tree in the initial posterior sample of trees we want to update with the new taxa.

This allows discarding branches that are unlikely candidates (i.e. low probability) for attaching the new taxon. The resulting attachment location is selected from a multinomial distribution with weights equal to the highest likelihood among the set of pendant branch lengths tested for each branch.

This heuristic could be refined at the cost of additional compute time. For example the likelihood profile of a fixed number of edges with the highest attachment probability can be improved by testing more attachment locations and additional potential pendant branch lengths (e.g. {0, median/2, median} instead of {0, median}). Alternatively, the posterior probability of attachment on each branch could be calculated directly (Matsen et al. 2010), however this may be too time consuming to be a practical improvement.

##### Parsimony 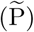 proposal

Alternatively, multinomial weights can be derived using parsimony scores, which are simply calculated using the first pass of the Fitch (1971) algorithm. The unnormalized attachment weight 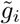 for the *i*th branch is calculated with the heuristic

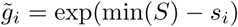

where *S* = (*s*_*i*_, …, *s*_2*n*−3_) is a vector containing the parsimony score of each attachment branch.

##### Heated parsimony (P) and likelihood (L) proposals

Through our simulations we noted that when simply normalizing the parsimony scores or likelihoods to a sampling distribution, the probability of the highest scoring branch is often several log units higher than the attachment probabilities of the other branches. Therefore the proposal often chooses the same branch with high probability. Unfortunately, for some sequences and tree configurations this attachment branch proposal algorithm can, with high probability, propose tree topologies that have low posterior support. That is, the attachment branch proposal distribution and posterior distribution can be poorly matched for some trees and sequences.

To mitigate the impact of poorly matching proposal and posterior distributions we explored a “heated” proposal distribution, created by raising the attachment probabilities to the power *α* = 0.05. This approach is inspired by the Metropolis-coupled MCMC method (Geyer 1991) in which the posterior distribution of a hot chain is exponentiated with a number less than one, hence flattening out the posterior distribution. The one-letter code of non-heated proposals is decorated with a tilde. For example, L̃ refers to the non-heated likelihood-based proposal and L denotes the corresponding heated proposal. In our implementation *α* was chosen by evaluating a range of possible values on an independent simulated test data set and is fixed throughout the simulations presented in the result section. Tuning the *α* parameter might improve the efficiency of the sampler. We have not explored this possibility.

#### Step 2: distal length choice

##### Uniform (U) proposal

The location on the attachment branch *e* to attach the pendant branch is drawn from a uniform distribution, 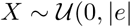, where 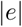 is the length of branch *e*.

##### Maximum likelihood normal (N) proposal

In this proposal scheme, the attachment location along branch *a* of the new branch and the pendant branch length are co-estimated using maximum likelihood. The distal length *x* is then drawn from a truncated normal distribution with location parameter *µ* equal to the maximum likelihood estimate (MLE) of the distal branch length. The distribution is truncated below 0 and above the length of the attachment branch. The scale parameter *σ* is arbitrarily chosen to be 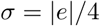. The distal length is set to 0 for 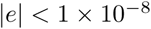.

##### Maximum likelihood asymptotic (A) proposal

This method proceeds in the same manner as N, except that the standard deviation in the proposal distribution is found using the posterior distribution around its maximum. Specifically, we use a quadratic approximation to the log likelihood distribution *ℒ* centered on the maximum likelihood estimate of *x*. That is,

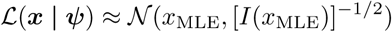

where *I*(*x*_MLE_) is the observed information

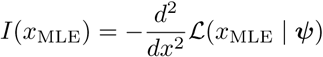

#### Step 3: branch length choice

##### Prior (P) proposal

The pendant branch length is simply drawn from the prior (e.g. *Y* ~ Exp(10)).

##### Maximum likelihood (M) proposal

The first guided method to draw the pendant branch length is similar in spirit to Step 2: the branch length is drawn from an exponential distribution with mean equal to the MLE of the pendant branch as calculated in the previous step.

##### Icfit (F) proposal

The second method makes use of a surrogate log-likelihood function to approximate the marginal posterior distribution of the pendant branch. This four-parameter surrogate function is specialized to the task of approximating single-branch phylogenetic likelihood functions. An implementation is available at https://github.com/matsengrp/lcfit and a manuscript describing it is in preparation (Claywell et al. 2017). For completeness we outline the method here.

Let *f* be the lcfit function, which is defined by four non-negative parameters, and evaluated at branch lengths *t:*

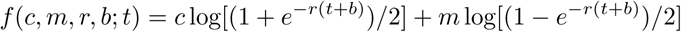

Ignoring the parameter *b*, this is the log-likelihood function of the binary substitution model on a two-taxon sequence alignment with *c* constant sites, m mutated sites, and substitution rate *r*. The parameter *b* effectively allows truncation of the left hand side of the curve, which enables modeling of likelihood curves with nonzero likelihood at *t* = 0.

One can fit the likelihood curve efficiently with access to the maximum likelihood branch length, an estimate of the second derivative at this location, and several other sampled points. Once the parameters of the lcfit surrogate are fit, one can use rejection sampling to obtain samples from the surrogate posterior formed by the product of the surrogate likelihood and the branch length prior.

##### Simulations

We generated 5 replicates of 10, 50, and 100 taxon trees under the birth-death process (λ = 6, *µ* = 2) using the R package TreeSim (Stadler 2010). Data sets will be labeled D*x*T*y* where the *x* is the replicate index and the *y* corresponds to the number of taxa (e.g. data set D1T10 is the first of five data sets containing 10 taxa). For each tree a nucleotide alignment with 1000 sites was simulated using the Jukes-Cantor substitution model (JC69) using `bppseqgen` from the Bio++ package (Guéguen et al. 2013). The posterior distribution of each phylogeny was approximated in two independent runs using MrBayes with three chains (i.e. 2 heated chains) for 300,000 iterations, ensuring an average standard deviation of split frequencies (ASDSF) below 0.01. A uniform prior on the topology and an exponential prior with mean 0.1 on branch lengths were specified. The chain was thinned down to 1,000 samples, of which the first 250 iterations were discarded.

For each data set, 1, 3, or 5 sequences were removed and the posterior distribution of each tree was approximated again using MrBayes, as described above.

The resulting posteriors were used as a starting point for inference with the our `sts` software (described below). The, 1, 3, or 5 removed sequences were sequentially added to MrBayes posterior samples using `sts` to approximate the full posterior. `sts` used the same priors as in the MrBayes analysis. We tested various numbers of particles in our SMC runs, each of which was a multiple of the number of trees in the original sample (in this case, 750). Define the *particle factor* to be the number of particles in the SMC divided by the number of trees in the original sample.

## Results

We implemented a prototype of Online Phylogenetic Sequential Monte Carlo (OPSMC) in an open source software package called `sts`. `sts` implements several different transition kernels for updating a phylogenetic posterior with new sequences. These transition kernels are described in detail in the Methods section, and include a uniform proposal and more sophisticated proposals that were developed with the aim of sampling updated trees more efficiently. As a prototype developed to test transition kernels rather than for practical use, the current `sts` software implements only the JC69 model and uses the stratified resampling technique (Kitagawa 1996) as implemented in SMCTC (Johansen 2009). The current `sts` implementation can update an existing posterior distribution produced by MrBayes with new sequences. `sts` is available from http://github.com/OnlinePhylo/sts.

We evaluated the sequence addition proposals on datasets consisting of 10, 50, and 100 taxa using a variety of proposal step combinations for the transition kernel. These proposal combinations are indicated using a three letter code as described above. For example, in the following results 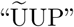 denotes a transition kernel constructed by proposing a branch uniformly at random (Step 1 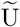), then proposing an attachment location uniformly along this branch (Step 2 U), and finally drawing a pendant branch length from the prior (Step 3 P).

In order to understand whether OPSMC is providing an accurate posterior approximation, we compare the OPSMC results after the addition of 1, 2, and 5 sequences to what was obtained by running MrBayes on the same datasets. In what follows we present the results based on data sets D1T50 and D1T100 in detail. Analysis of the other data sets showed similar results and these are provided as supplementary data.

### Effective Sample Size (ESS) from OPSMC

The guided proposals showed clearly superior ESS compared to the uniform proposal with any particle factor (Fig. 2). The ESS produced by guided proposals also shows a strong linear relationship with the number of particles. This relationship was predicted by Dinh et al. (2016), where the ESS of the sampler was bounded below by a constant multiple of the number of particles. These linear regressions have different slopes, suggesting that as the user targets higher ESSs the differences between proposals become more important. In Figure 2 we find that proposals using likelihood to select an attachment branch (Step 1 L) have higher ESS than parsimony, while use of lcfit to propose pendant branch lengths (Step 3 F) yields a large advantage in ESS.

**Figure 2.**
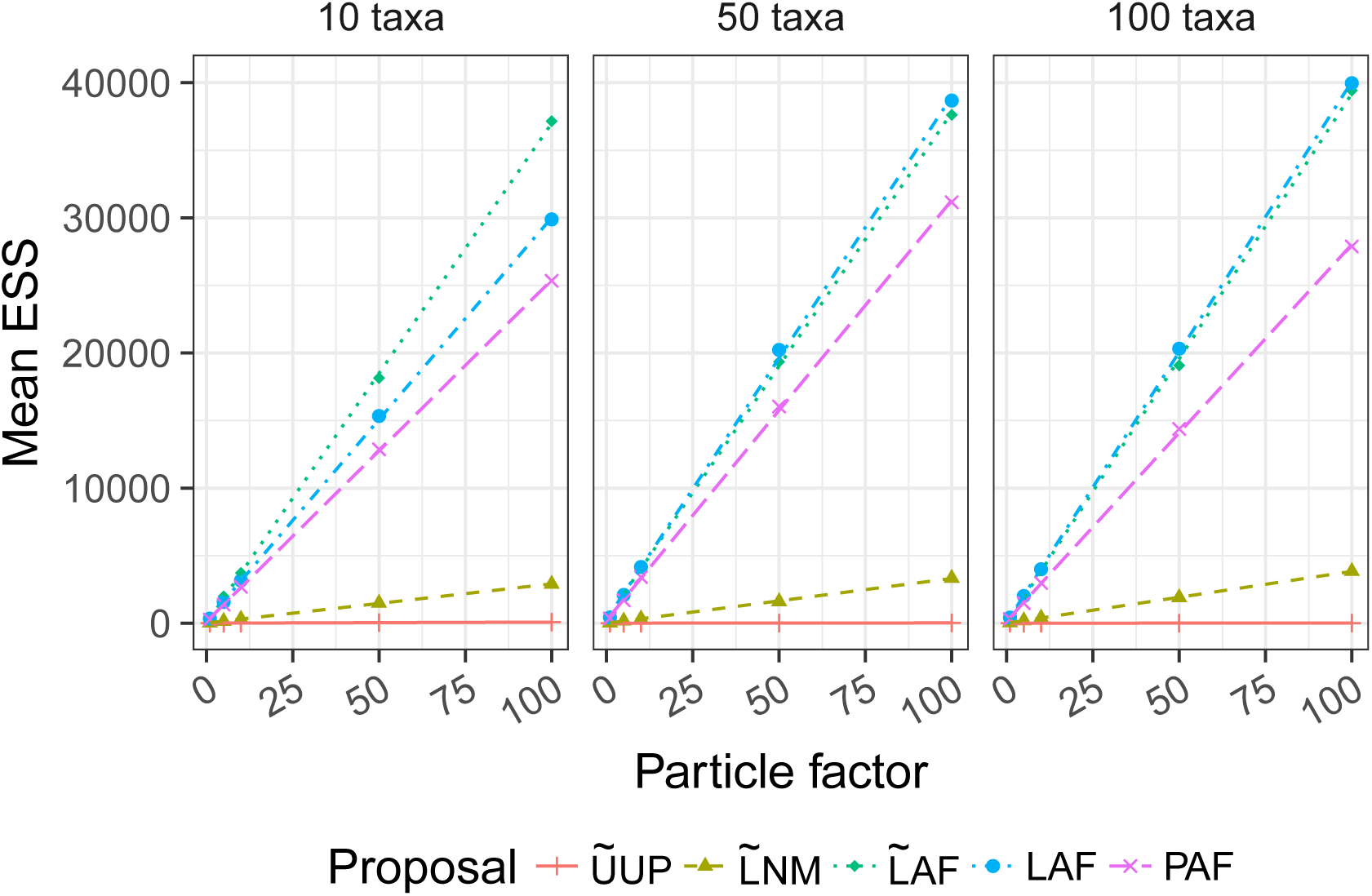
Mean effective sample size (ESS) as a function of the particle factor across every data set.

As discussed further below, we unfortunately cannot always equate high ESS with good quality posterior samples since ESS alone does not guarantee the OPSMC is sampling a high probability region. Therefore we performed a detailed comparison with the MrBayes posteriors.

### Comparison of posteriors from SMC and MrBayes MCMC

We measured the distance between the maximum likelihood tree inferred with PhyML (Guindon et al. 2010) and each tree in the MrBayes posterior samples and the OPSMC posterior samples using the weighted Robinson-Foulds (L1-norm) distance (Robinson and Foulds 1981) calculated with the DendroPy library (Sukumaran and Holder 2010). Under the assumption that the MCMC implemented in MrBayes has been run long enough to accurately approximate the true posterior, the ability of each OPSMC proposal scheme to approximate the true posterior distribution can be assessed by comparing the distribution of their L1-norm distances to that produced by MrBayes.

The results indicate that guided proposals, especially PAF and LAF, yield superior posterior approximations to those produced by UUP with the particle counts used here (Figs. 3, 4).

**Figure 3.**
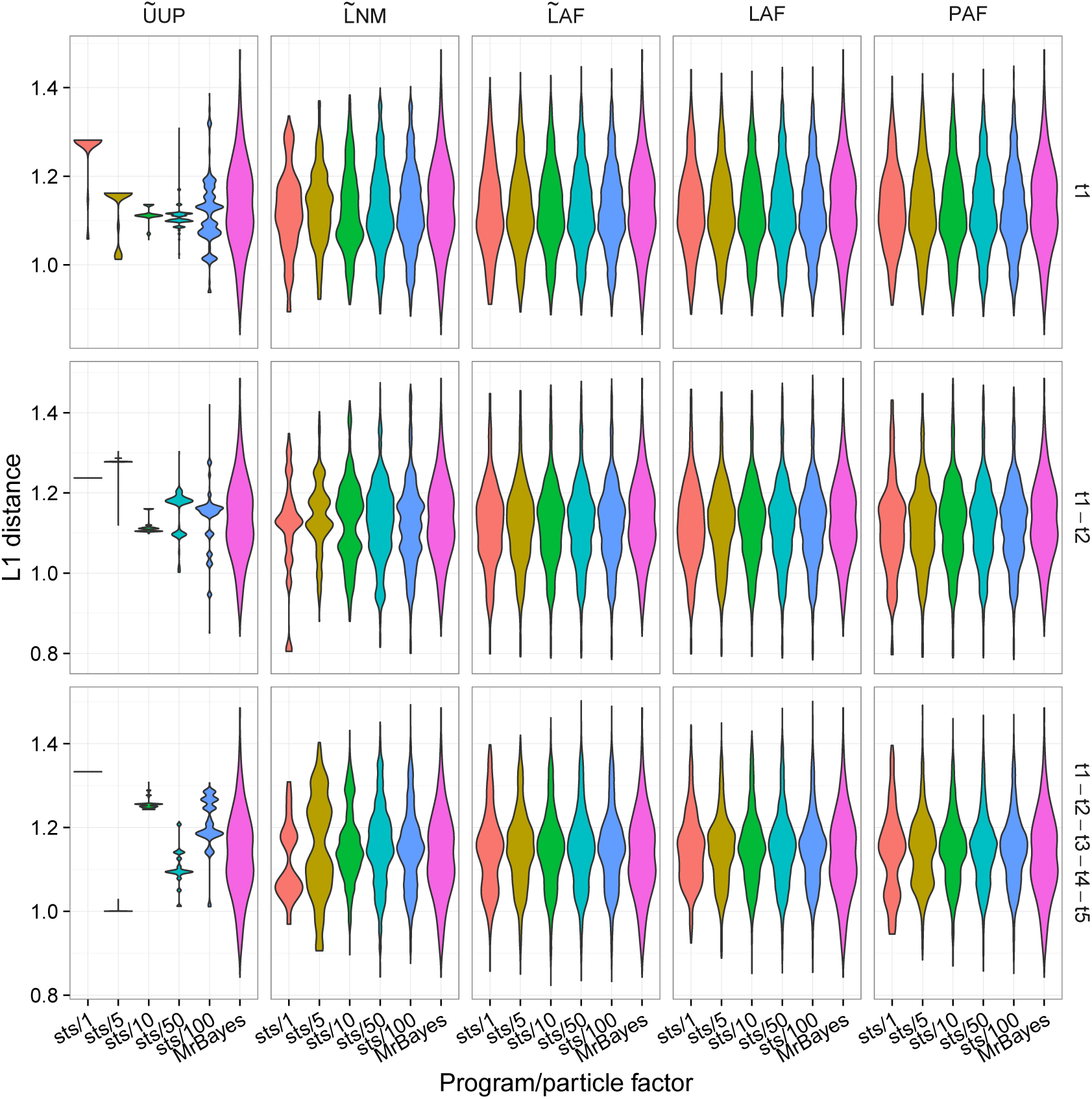
Posterior distribution of weighted Robinson-Foulds distances between each tree generated by `sts` and the maximum likelihood tree inferred by PhyML. These results are for data set D1T50. Labels on the right of the y axis indicate which taxa were removed.

**Figure 4.**
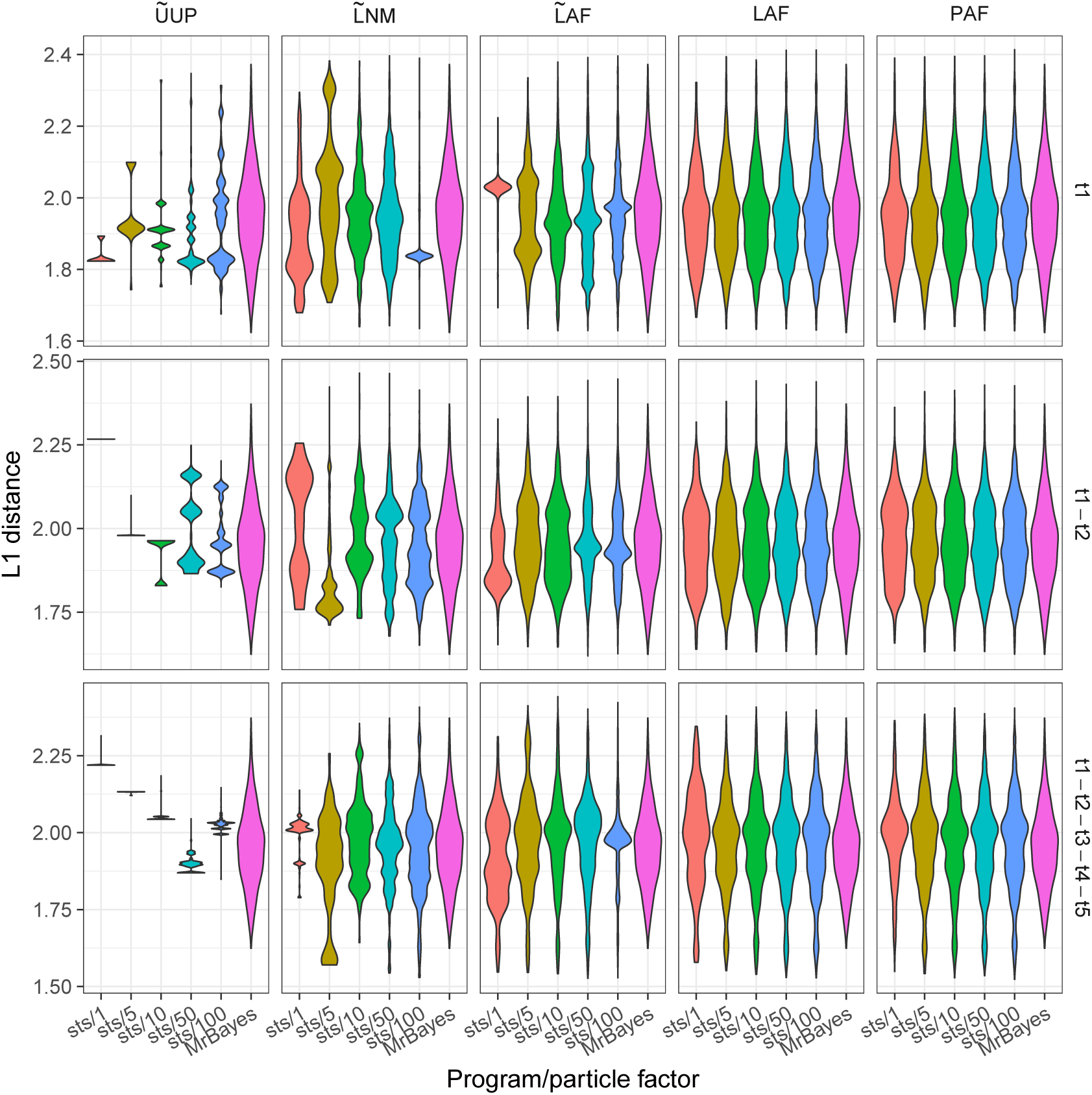
Posterior distribution of weighted Robinson-Foulds distances between each tree generated by `sts` and the maximum likelihood tree inferred by PhyML. These results are for data set D1T100. Labels on the right of the y axis indicate which taxa were removed.

### Measuring convergence with the Average Standard Deviation of Split Frequencies (ASDSF)

The average standard deviation of split frequencies (ASDSF) (Lakner et al. 2008; Ronquist et al. 2012) is a widely employed statistic used to assess the convergence of independent MCMC analyses. The ASDSF approaches zero when the set of topologies contained in the posterior approximations of different Monte Carlo sampling runs have converged. We used this metric to determine whether the posterior approximation produced by sequential taxon addition in `sts` is similar to that which would have been produced by simply running MrBayes on the complete data set. We calculated the ASDSF between the updated posterior distribution generated by `sts` and an independent MrBayes analysis on the full data set. It is common practice to use ASDSF as a convergence criterion, stopping Bayesian MCMC once the ASDSF is less than 0.01 (Lakner et al. 2008).

We find that that guided proposals can yield an ASDSF less than 0.01, even with relatively small particle factors (Fig. 5). In contrast, unguided proposals such as 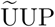 consistently yielded posterior approximations with an ASDSF that was an order of magnitude higher (worse), even when large particle systems were employed (particle factor 100, with 75000 particles). Also, the simple likelihood-based guided proposals (Step 1 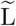) fail to yield posterior approximations that meet the convergence criterion, whereas their heated relatives perform much better.

**Figure 5.**
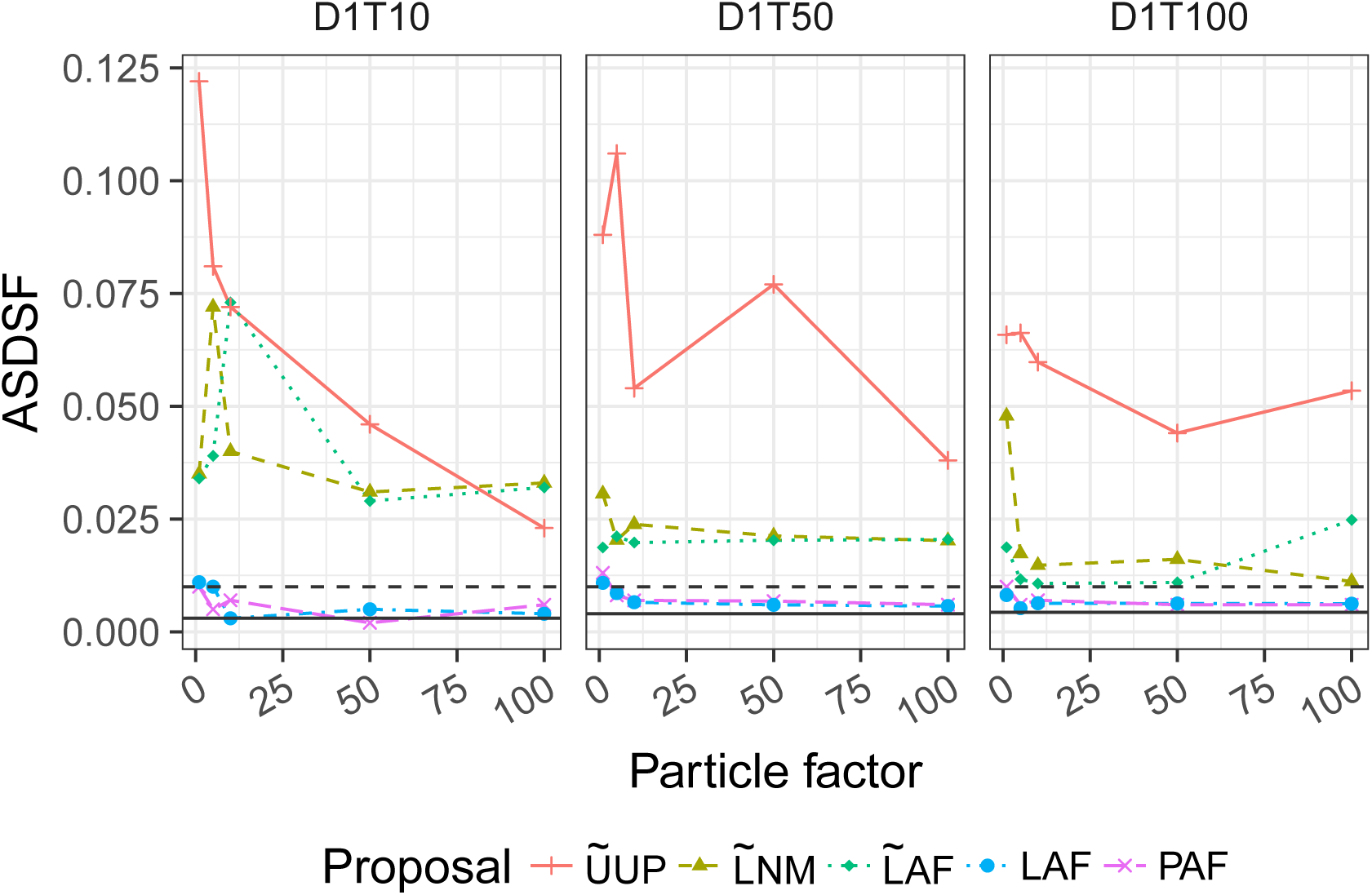
Average standard deviation of split frequencies (ASDSF) between `sts` and MrBayes posterior samples. These results are for the three data sets D1T10, D1T50, D1T100 and include several particle factors. The horizontal line represents the ASDSF calculated using two chains in MrBayes. The horizontal dashed line marks 0.01, a common convergence criterion.

### Compute time of OPSMC proposal schemes

In previous studies of phylogenetic sequential Monte Carlo (Bouchard-Côté et al. 2012; Wang et al. 2015), the number of peeling recurrences were used as a proxy to compare the running time of different proposals. Since some non-likelihood aspects of our implementation (such as calculation of the parsimony score in the PAF proposal) incur a non-negligible compute load, we directly investigated the wall clock time for each proposal instead.

As expected, we find that uniform proposals are at least an order of magnitude faster per proposal operation than the guided proposals (Supplementary Figs. S4,S5). The timing results also suggest that in the current implementation, the `lcfit` approximation (Step 3 F) incurs a significant cost in compute time relative to the other approaches.

However, when measured in terms of compute time required per unit of ESS in the resulting sample, we find that the guided proposals outperform 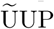 by a large margin (Fig. 6). We note that in SMC, a high ESS is necessary (but not sufficient) for an accurate posterior approximation. Interestingly, the results show that the extra compute time used by the normal approximation in step 2 and lcfit in step 3 may be justified since the 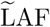 proposal has on average superior ESS per unit time relative to 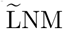. For some replicates (e.g. D5T10) the runtime-to-ESS ratio is much higher for the 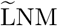 and 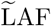 proposals than for the heated proposals. The ESSs of those runs are extremely low (Supplementary Fig. S2), highlighting that some data sets are more difficult to analyze.

**Figure 6.**
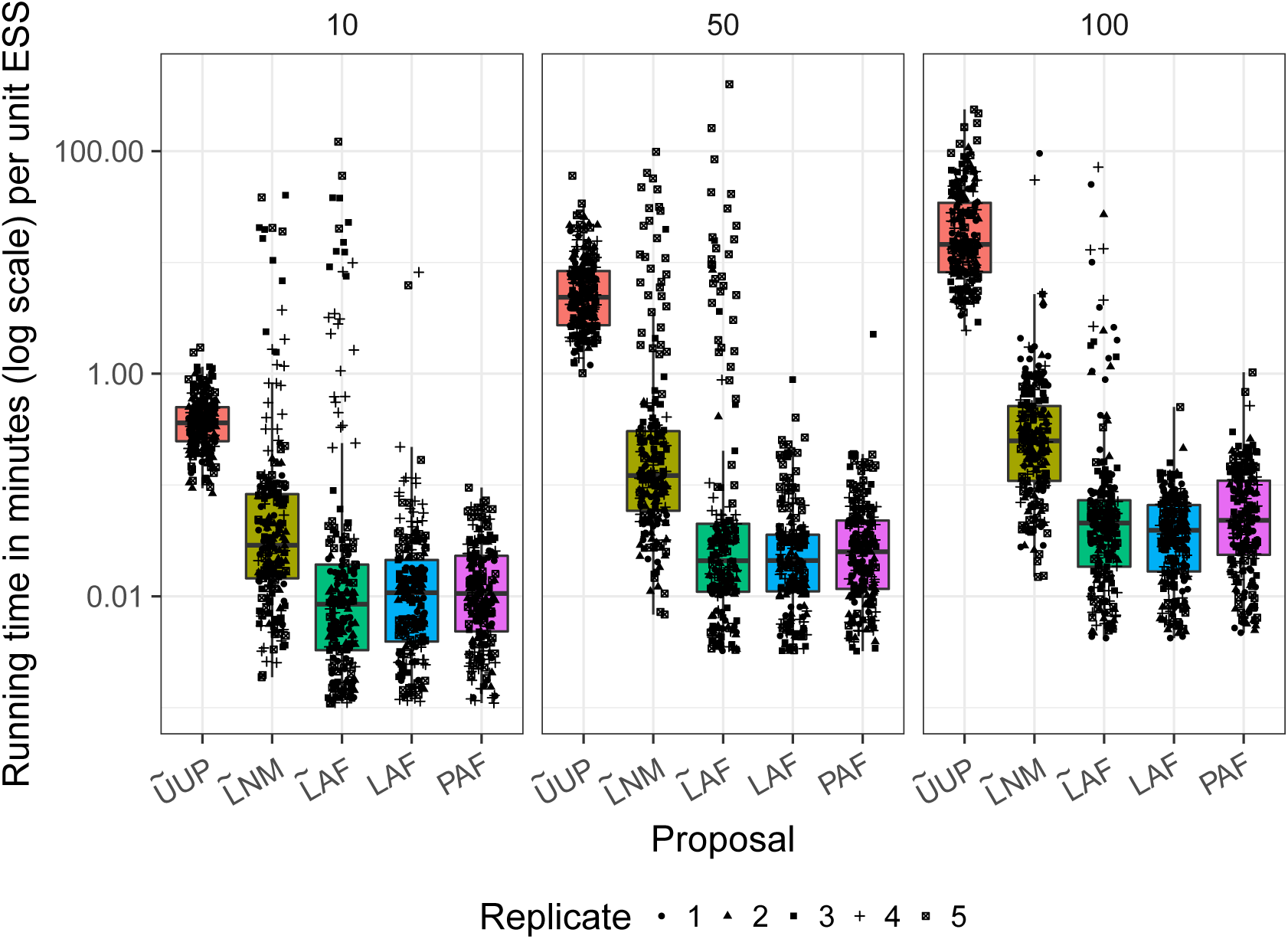
Running time in minutes per effective sample size (ESS) unit for each proposal method. These results are for every data set and include all five particle factors. Numerical labels on top of each subplot give the number of taxa in each data set.

Next, we evaluated compute time in a situation that commonly arises in genomic epidemiology, where an updated phylogenetic posterior is desired every time a new sequence becomes available. We thus simulated the sequential arrival of five taxa, comparing the time required for `sts` to update posteriors against the approach of sampling each of the five posteriors from scratch with MCMC. These new sequences are added to an existing data set containing 45 or 95 sequences. In an offline setting, this is equivalent to running MrBayes once each on alignments containing 46 to 50 or 96 to 100 sequences. We compared the total amount of time for the five MrBayes runs to a single run of `sts` with the same five sequences. For each method, we report the minimum time required to compute a posterior approximation that meets the widely used Monte Carlo convergence criterion of an ASDSF lower than 0.01 (Fig. 7). The plot shows the results based on ten data sets (D1T50-D5T50 and D1T100-D5T100) that we described above and for each replicate the five sequences were sequentially added in three different orders across three runs. Runs from `sts` that do not reach an ASDSF below 0.01 are not included in the plots, therefore each panel contains at most 15 points for `sts`. We find that `sts` is faster than MrBayes and that PAF and LAF required a particle factor of only one in 11 and 14 cases, respectively, for data sets containing 50 sequences. LAF performed marginally better than PAF for the larger 100 taxon data set, wherein the LAF proposal reached the target ASDSF 12 times while PAF only reached it 3 times (See Table S1). In contrast, UUP was not able to sample trees that would meet the convergence criterion.

**Figure 7.**
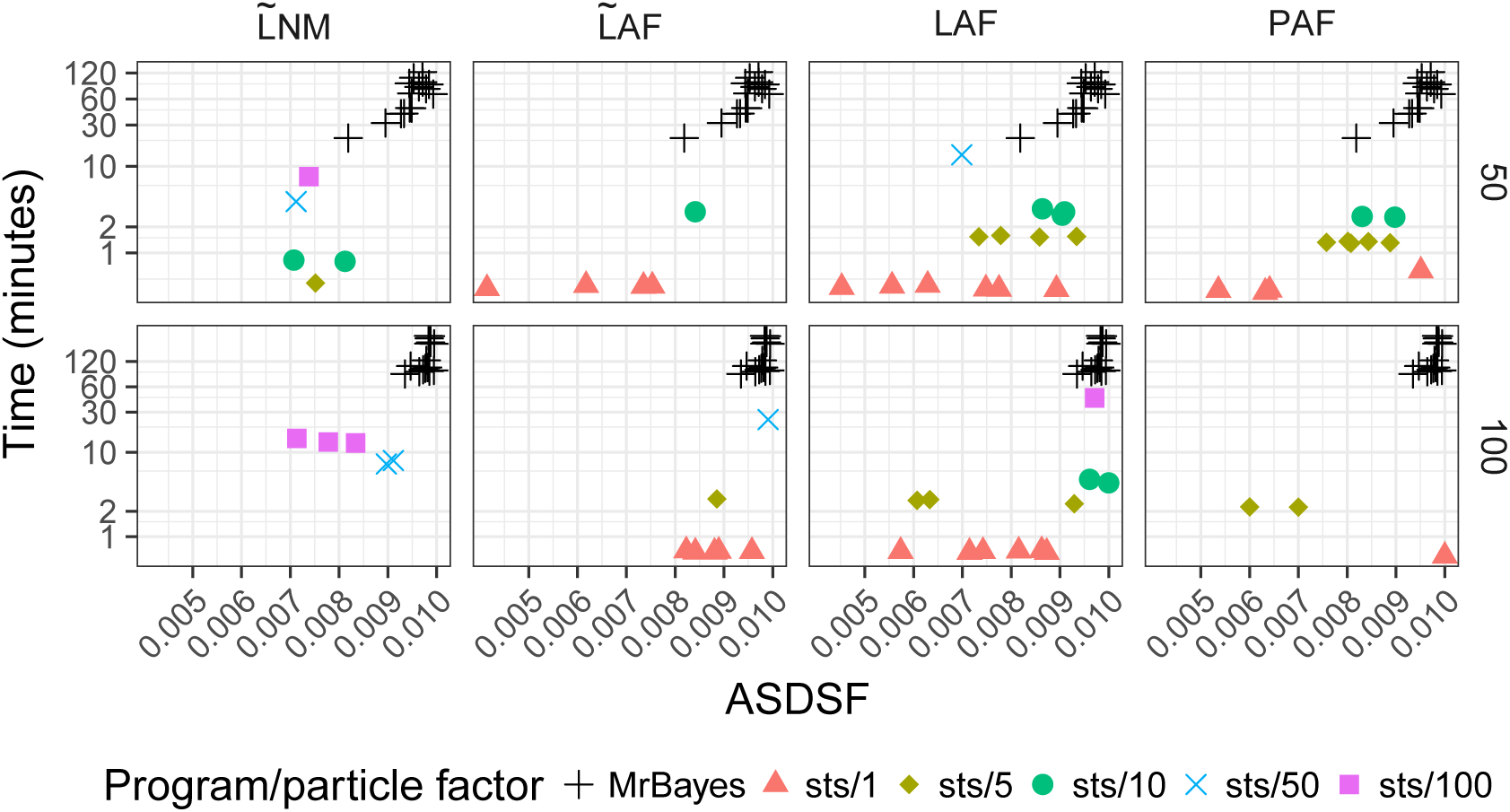
Running time in minutes (on a log scale) to sequentially add five new sequences to a data set of 45 or 95 sequences with MrBayes and `sts`. Here `sts` is run with particle factors ranging from 1 to 100 and the lowest time reported that achieved an ASDSF less than 0.010. Runs for which no particle factors resulted in an ASDSF less than 0.010 are not shown on the plot. Results are based on five data sets in which the taxa were added in three different orders, resulting in at most 15 data points. For example, the LAF plots contain 14 and 12 data points for data sets consisting of 50 and 100 sequences respectively (See supplementary table 1 for a full description of the other proposals).

The other non-heated methods provide intermediate results. We note that opportunities exist for further reduction of compute time required for `sts` (see Discussion), and that more significant efforts have been made to optimise MrBayes 3.2 (Ronquist et al. 2012), so the results here could be interpreted as a rough lower bound for the speed advantage that OPSMC may have over sequential runs of MCMC.

## Discussion

Here we have implemented the Online Phylogenetic Sequential Monte Carlo (OPSMC) framework described by Dinh et al. (2016) and evaluated how several alternative proposal schemes behave on synthetic data sets.

### Related work

Everitt et al. (2016) have also developed theory and an implementation for SMC on phylogenies. In their case they are focused on inferring ultrametric trees in a coalescent framework, whereas OPSMC as described here is for unrooted trees. Their clever attachment proposal is described in terms of lineage (path from tip to leaf) and branching time. They use proposals choosing lineage based on differences from the leaf to be attached and the existing leaves using a distribution based on Ewens’ sampling formula (Ewens 1972), and a branching time which also uses pairwise differences. They also make an interesting suggestion to ease the transition between the different posterior distributions by using “intermediate distributions.” However, they do not compare their output to samples from an existing MCMC phylogenetics package and they have not yet provided an open source implementation that would allow others to do so.

### Guided proposals work

In contrast with previous suggestions (Bouchard-Côté et al. 2012), we show that guided proposal schemes can greatly improve both the computational efficiency and the accuracy of a phylogenetic posterior approximation over simple uniform proposal schemes. When quality of posterior approximation is measured by either L1-norm (Figs. 3, 4) or ASDSF (Fig. 5), the LAF and PAF proposals clearly outperform the other schemes. Both LAF and PAF are able to achieve ASDSF below the 0.01 threshold that is typically used as an indicator of MCMC convergence, and can do so even with relatively small particle system sizes. This finding is especially important for the future application of SMC to phylogenetics, suggesting that proposal efficiency matters much more for SMC than MCMC.

### High ESS does not imply accurate posterior approximation

On the other hand the 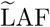 proposal provides the highest ESSs among any of the various proposal schemes, yet fails to achieve a low ASDSF, suggesting a poor posterior approximation. Detailed inspection of the behavior of 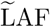 reveals that, without heating the likelihoods, the highest scoring attachment branch can be several log units above the others (including the correct branch), resulting in a multinomial weight close to 1 while the others are close to zero. The bias leads 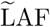 to always propose the same attachment branch even when a large particle system is employed, causing it to sample a narrow region of the tree space. The subsequent Step 2 and 3 proposals yield a large number of configurations with similar weights, resulting in a high ESS.

### Opportunities for computational optimisation

Particle degeneracy is a well known drawback of SMC algorithms, characterized by a large number of identical particles after the resampling step. However, particle degeneracy offers an opportunity for computational optimisation of guided proposals. The Step 1 proposal distributions for identical particles can be computed once and then reused. Similarly, Step 2 involves repeatedly calculating maximum likelihood estimates of the distal and pendant branch lengths, which are identical for identical trees. For the cost of a modest amount of bookkeeping, those estimates can be computed just once for each tree in the particle system, rather than once for each particle, yielding a significant speedup.

### Limitations

#### Limitations: OPSMC proposals

A common feature of the proposals we evaluated are that they consist of 3 steps: (1) selection of an attachment branch, (2) selection of an attachment point on the attachment branch, and (3) selection of a pendant branch length. However, the three step structure imposes some potentially undesirable restrictions on the resulting trees. One such restriction is that the length of the attachment branch selected in Step 1 is fixed, and is not adjusted in steps 2 or 3. Another potential issue is that the length of two adjacent branches in a phylogenetic tree can be strongly correlated. The current 3 step proposal scheme does not account for this correlation structure. The efficiency of the sampler could in principle be increased by combining the proposals in steps 2 and 3 to account for the dependency in lengths of the three branches incident to the attachment point. The branch lengths could be modeled as a multivariate truncated normal distribution where the covariance matrix captures correlations across branches.

Alternatively, an extension of the surrogate function described in the lcfit algorithm to multiple branches could improve the proposal.

#### Limitations: computational complexity

The computational complexity of choosing the attachment branch (i.e. Step 1) grows linearly with the number of taxa in the tree. Profiling of our implementation indicates that at the data set sizes we evaluated, Step 1 consumes less than 5% of total compute time. Nevertheless, scaling our approach to thousands of sequences or beyond is likely to require new heuristics or other techniques to reduce the time complexity of branch selection in Step 1. This is in contrast with drawing branch lengths in the other steps, which has a constant complexity with respect to the number of taxa. As in other standard SMCs, the memory requirement of `sts` scales linearly with the number of particles. The development of memory-efficient (Jun and Bouchard-Côté 2014) and highly-parallel (Paige et al. 2014) variants would be an essential step for scaling to large data sets.

#### Limitations: evolutionary models

The present work has focused on updating the tree topology and branch lengths in the very simple JC69 model of sequence evolution. In practice, richer phylogenetic models will almost always be preferred as they can provide a better fit to the sequence data, for example by modeling unequal rates of nucleotide substitution or clade-specific evolutionary rates. These additional model parameters are often continuous real-valued parameters. One way to sample these parameters was described and implemented by Bouchard and colleagues (Wang et al. 2015) who developed a method based on particle MCMC (Andrieu et al. 2010). Particle MCMC uses SMC on the tree topology and branch lengths to approximate the marginal likelihood of the remaining continuous parameters, which are sampled using MCMC moves. A similar particle MCMC approach, or another means to sample the evolutionary model parameters has yet to be developed in the context of online phylogenetic SMC.

#### Limitations: path degeneracy

OPSMC, like all SMC algorithms, is prone to path degeneracy, especially when the sampler iterates through generations with low ESS. As previously suggested (Dinh et al. 2016; Bouchard-Côté et al. 2012), the use of MCMC moves between iterations of an SMC can help alleviate the path degeneracy problem. Although some simple MCMC moves have been implemented in `sts`, preliminary results suggest that such a large number of these simple moves would be required to address the path degeneracy problem that a better result can be achieved by simply using a larger particle system. Dinh et al. (2016) suggest a valid sampler could be constructed using any mix of MCMC and SMC moves, ranging from entirely SMC to almost entirely MCMC, but a thorough investigation of the optimal blend, incorporating known advanced MCMC proposals (Ronquist et al. 2012), is yet to be done.

#### Limitations: convergence diagnostics

In the current work we have evaluated the accuracy of the OPSMC’s posterior approximation by comparing the sample to a collection of MCMC samples derived from MrBayes. This approach is inviable in practice, since an independent posterior approximation derived from MCMC will not generally be available. Further work is needed to develop and evaluate comparable approaches for convergence diagnostics to be used with OPSMC.

### Conclusion

Phylogenetic inference is quickly becoming an essential tool in modern infectious disease epidemiology. When sequence data arrives continuously, as in the case of an outbreak, it would be preferable to simply update a previous analysis rather than recomputing results for all sequences. We have presented the first investigation into the practical feasibility of online Bayesian phylogenetics using Sequential Monte Carlo. Our findings suggest that the choice of proposal distribution is especially important for successful inference with OPSMC, and to this end we have described several transition kernels and evaluated their strengths and weaknesses. We have also found that simple likelihood-based proposals can strongly bias the proposal distribution away from the posterior, and have shown that smoothing of these proposals can yield a posterior approximation that meets the *de facto* standard criteria for topological convergence in phylogenetic MCMC. The current `sts` implementation is limited to the simplest evolutionary models, and although our initial findings are promising, significant future work will be required to integrate the approach into the familiar software packages that implement the more complex evolutionary models in common use today.

## Acknowledgements

The authors would like to thank Aaron Gallagher for his early contributions to the `sts` source code, and to Julien Dutheil and Bastien Boussau for assistance with the Bio++ suite. VD and FAM funded by National Science Foundation grants DMS-1223057, CISE-1564137, and National Institutes of Health grant U54-GM111274. FAM supported by a Faculty Scholar grant from the Howard Hughes Medical Institute and the Simons Foundation.

## SUPPLEMENTARY MATERIAL

**Table S1.**
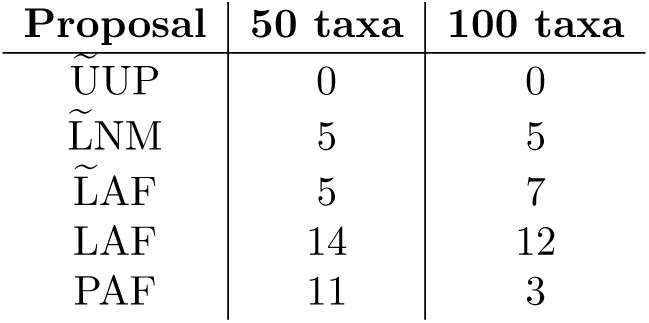
A summary of the number of `sts` runs (out of 15 total) that that reached an average standard deviation of split frequency (ASDSF) of less than 0.01.

| Proposal | 50 taxa | 100 taxa |
| --- | --- | --- |
| $\tilde{\text{UUP}}$ | 0 | 0 |
| $\tilde{\text{LNM}}$ | 5 | 5 |
| $\tilde{\text{LAF}}$ | 5 | 7 |
| LAF | 14 | 12 |
| PAF | 11 | 3 |

**Figure S1.**
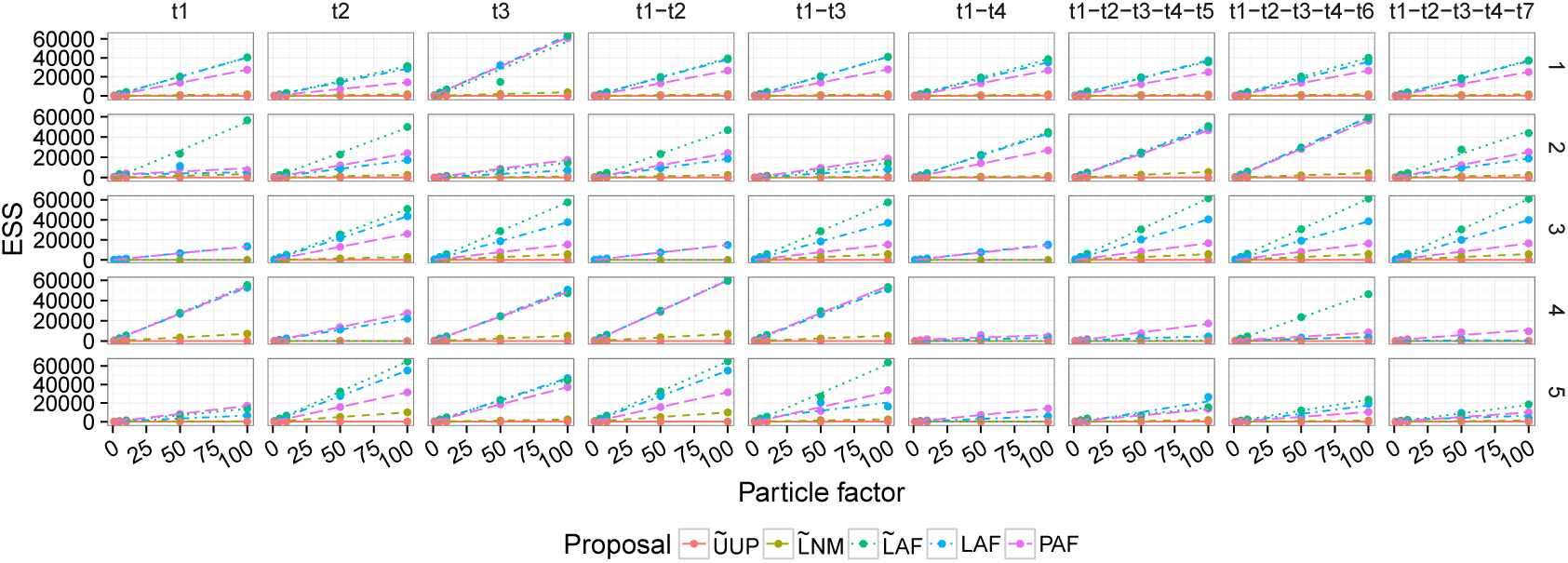
Effective sample size (ESS) as a function of the particle factor across every data set of size 10. Labels on top of each subplot identify the taxa that were removed. For subplots labeled *t1-t2*, taxa *t1* and *t2* were removed.

**Figure S2.**
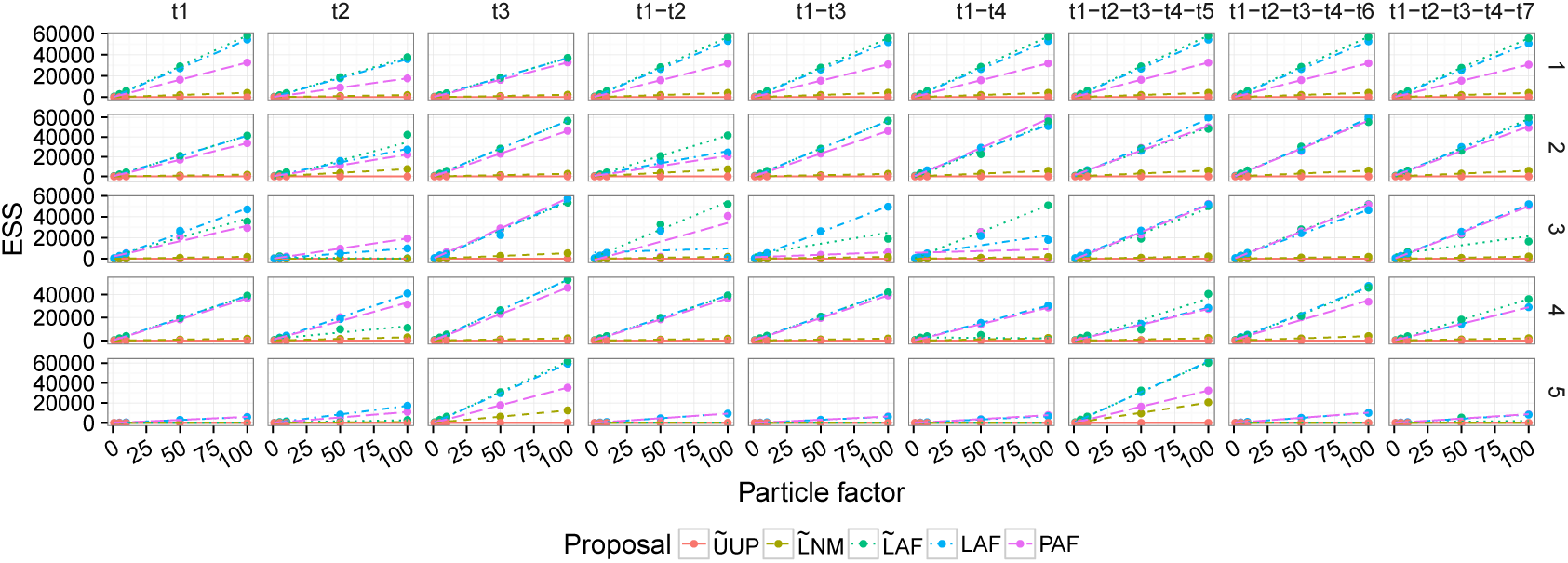
Effective sample size (ESS) as a function of the particle factor across every data set of size 50. Labels on top of each subplot identify the taxa that were removed. For subplots labeled *t1-t2*, taxa *t1* and *t2* were removed.

**Figure S3.**
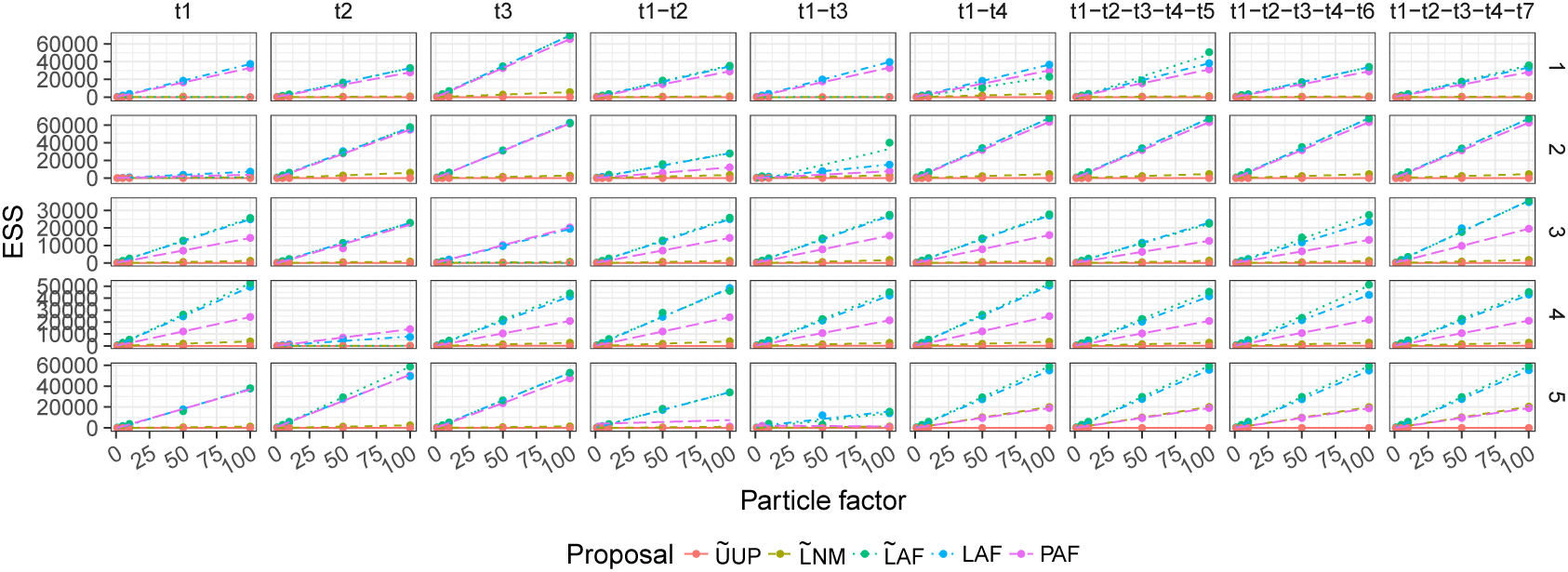
Effective sample size (ESS) as a function of the particle factor across every data set of size 100. Labels on top of each subplot identify the taxa that were removed. For subplots labeled *t1-t2,* taxa *t1* and *t2* were removed.

**Figure S4.**
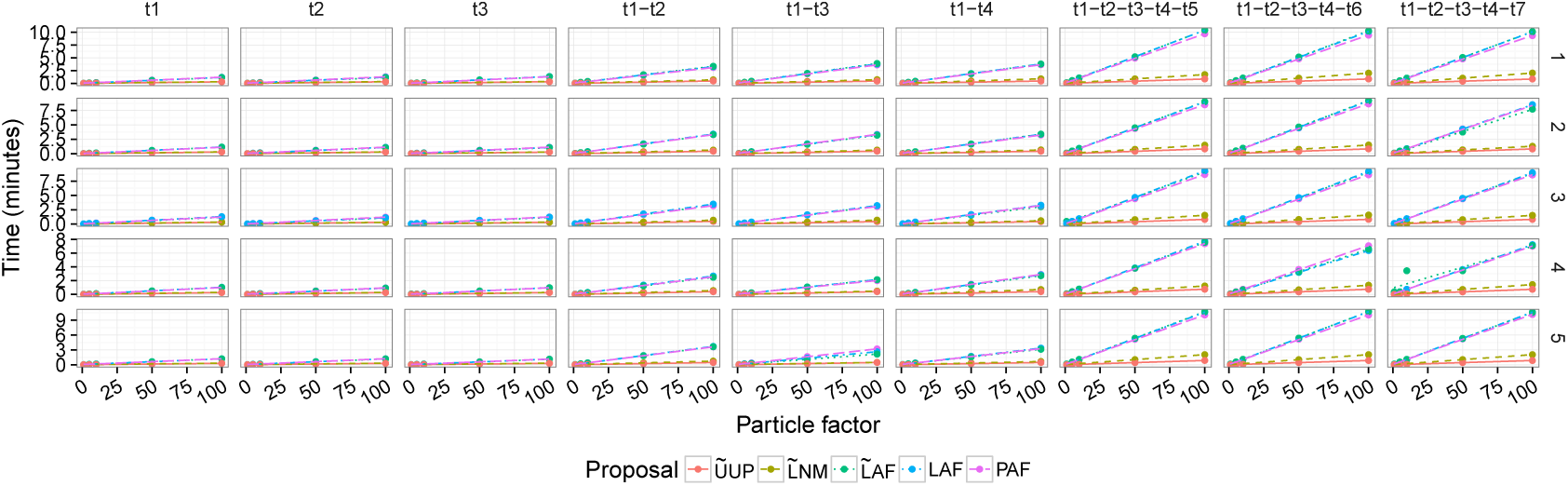
Running time in minutes per effective sample size (ESS) unit for each proposal method. across every data set of size 10.

**Figure S5.**
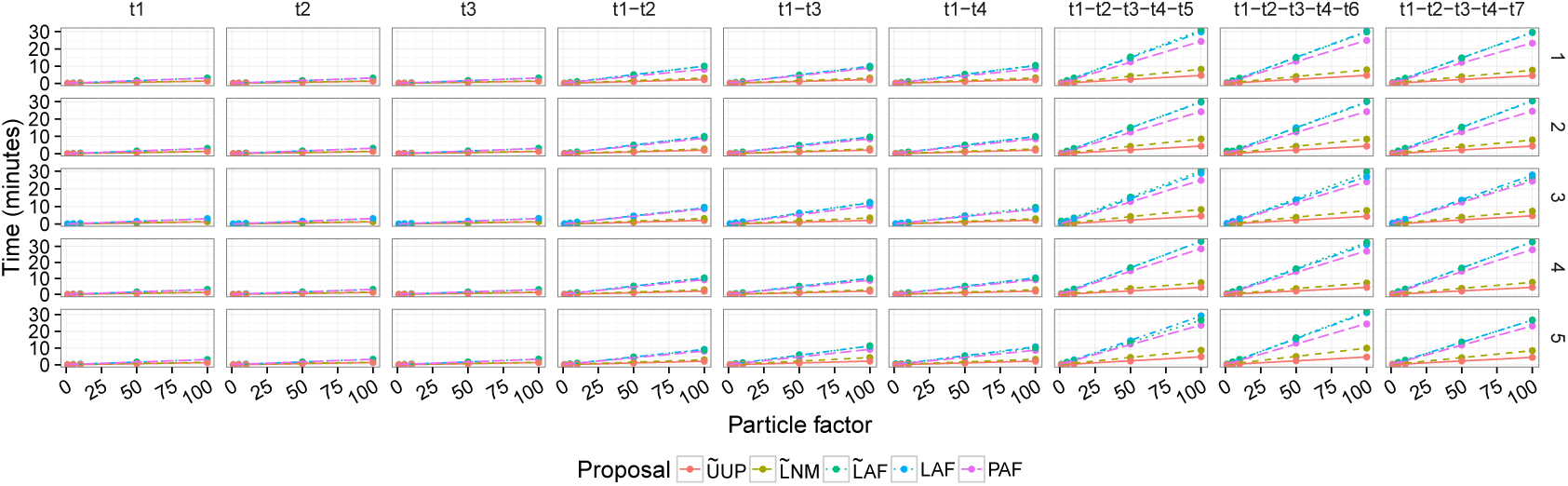
Running time in minutes per effective sample size (ESS) unit for each proposal method. across every data set of size 50.

**Figure S6.**
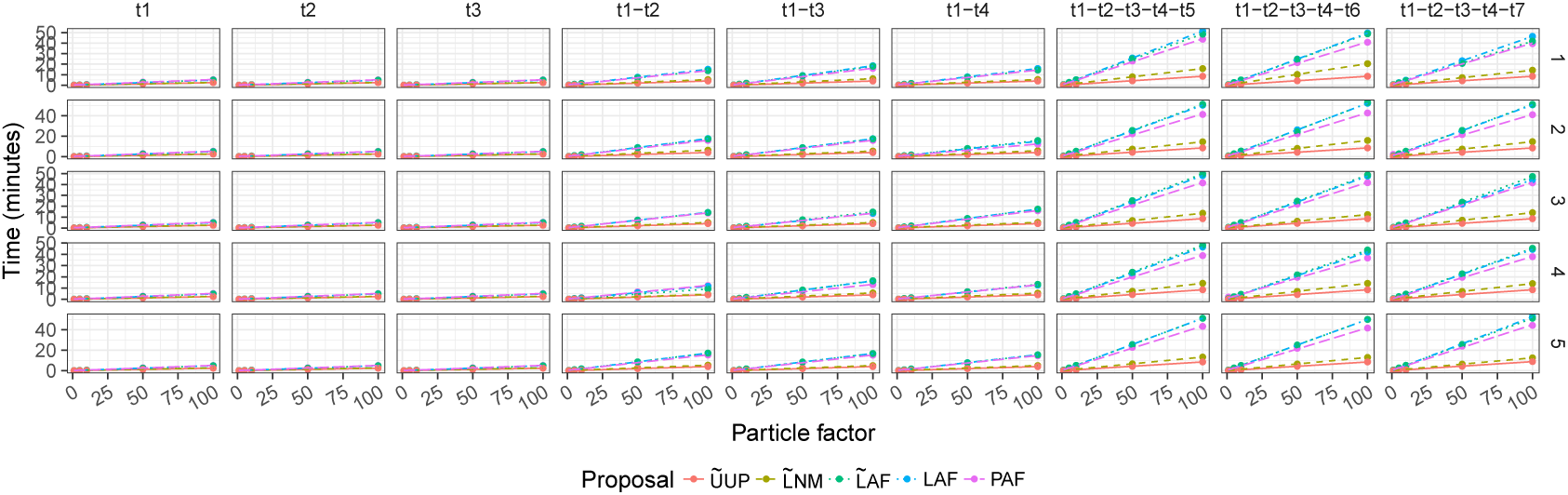
Running time in minutes per effective sample size (ESS) unit for each proposal method. across every data set of size 100.

**Figure S7.**
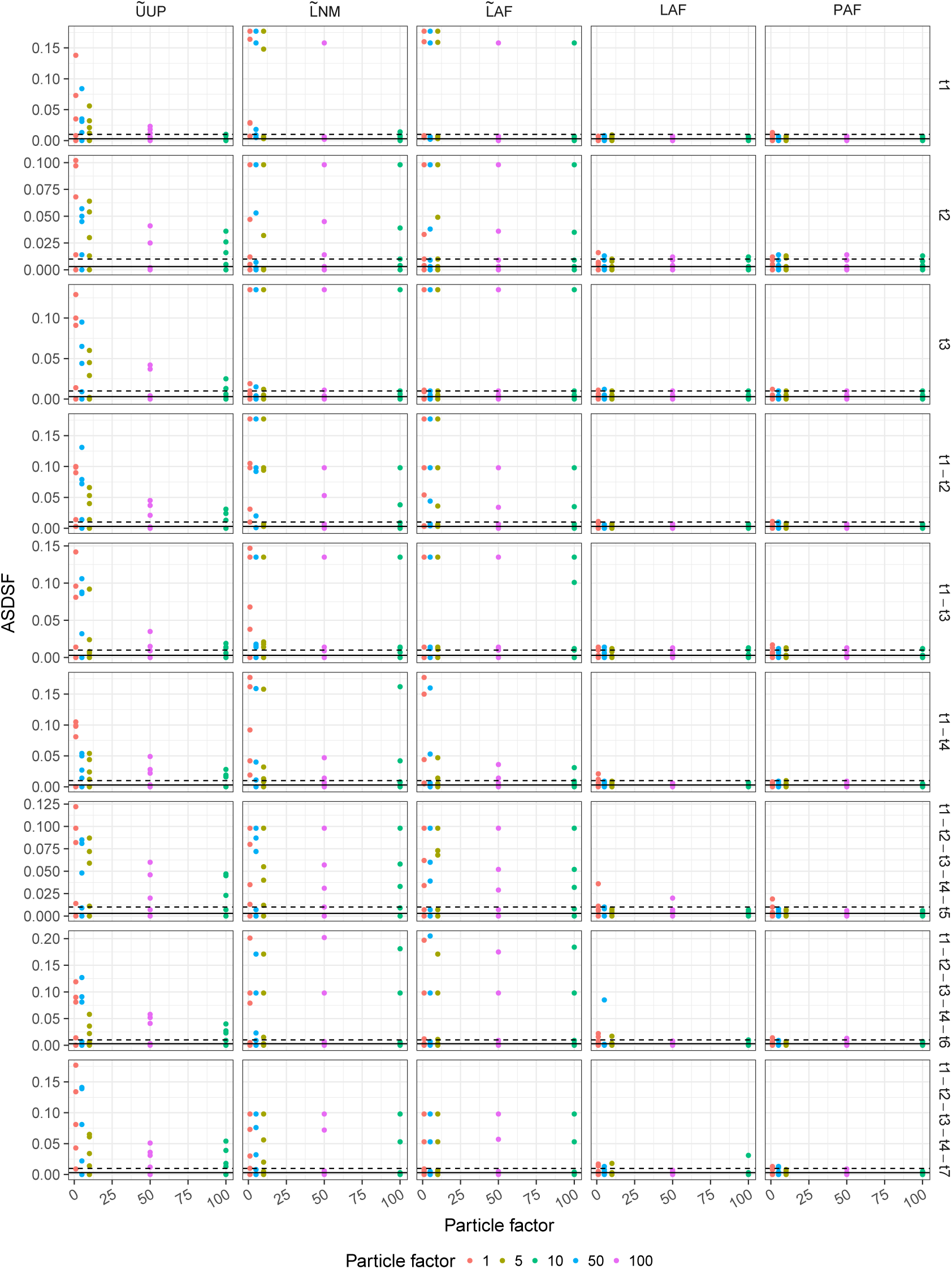
Average standard deviation of split frequencies (ASDSF) as a function of the particle factor across every data set of size 10.

**Figure S8.**
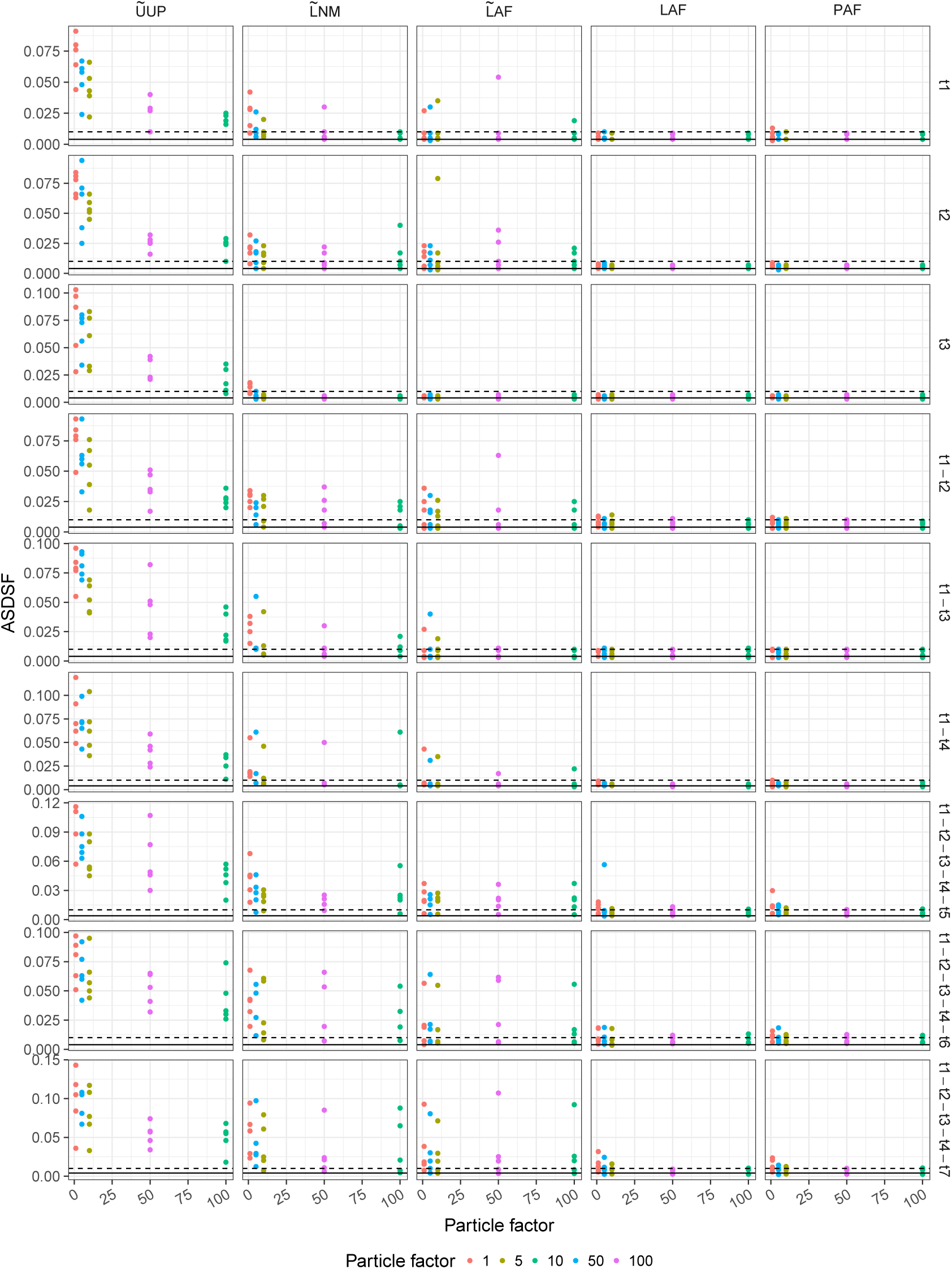
Average standard deviation of split frequencies (ASDSF) as a function of the particle factor across every data set of size 50.

**Figure S9.**
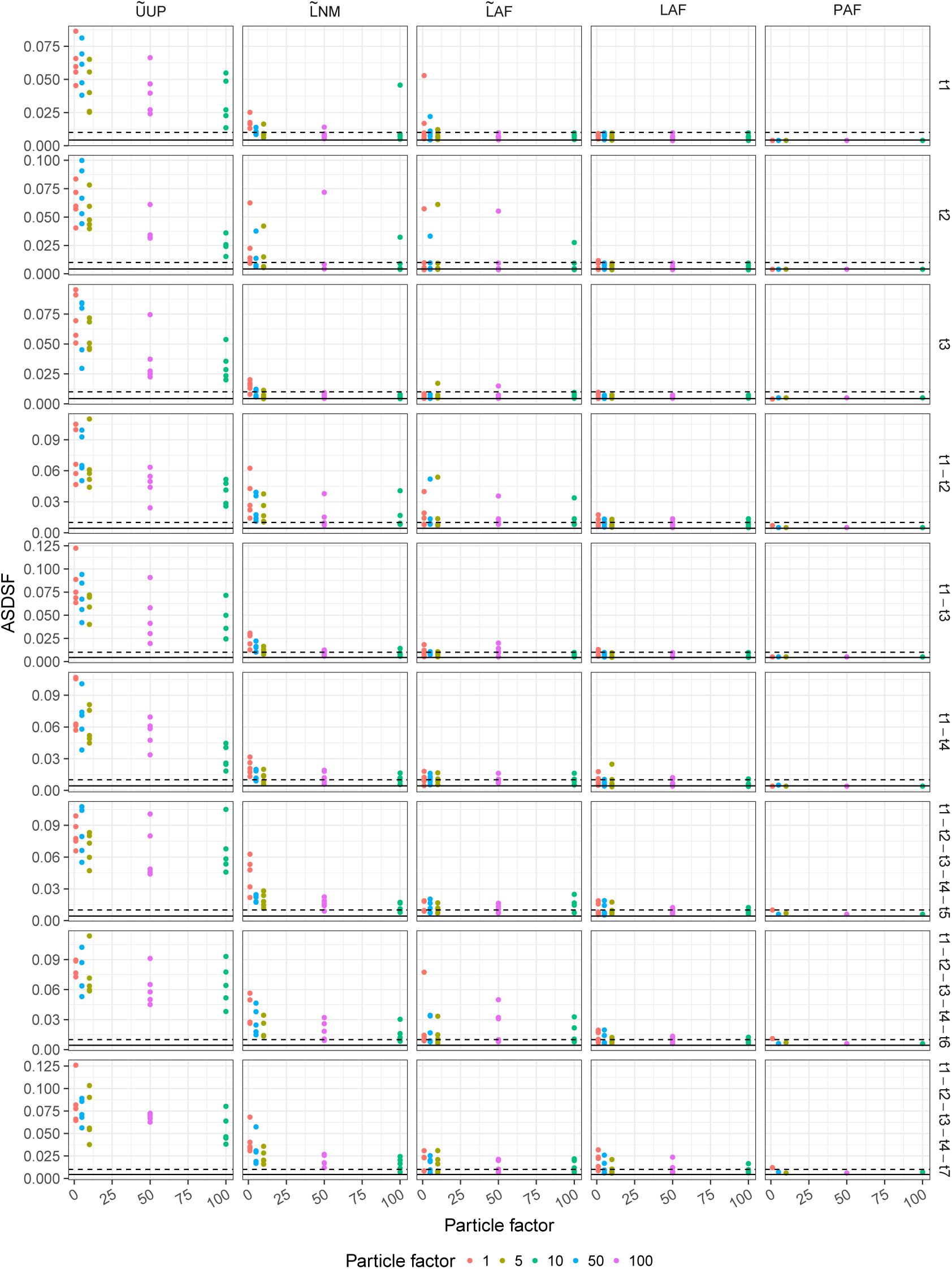
Average standard deviation of split frequencies (ASDSF) as a function of the particle factor across every data set of size 100.

**Figure S10.**
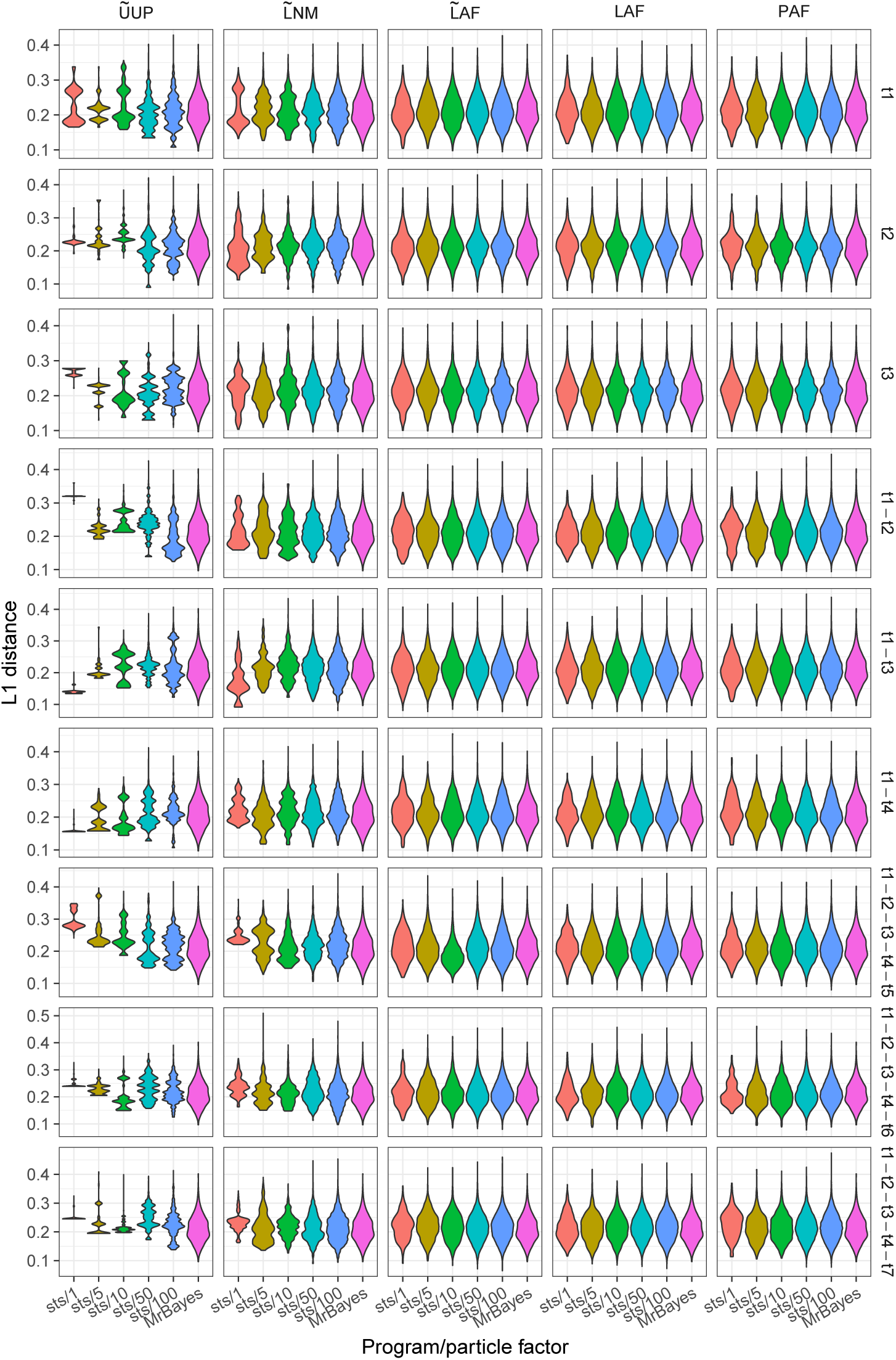
Posterior distribution of weighted Robinson-Foulds distances between each tree generated by `sts`and the maximum likelihood tree inferred by PhyML using the D1T10 data set.

**Figure S11.**
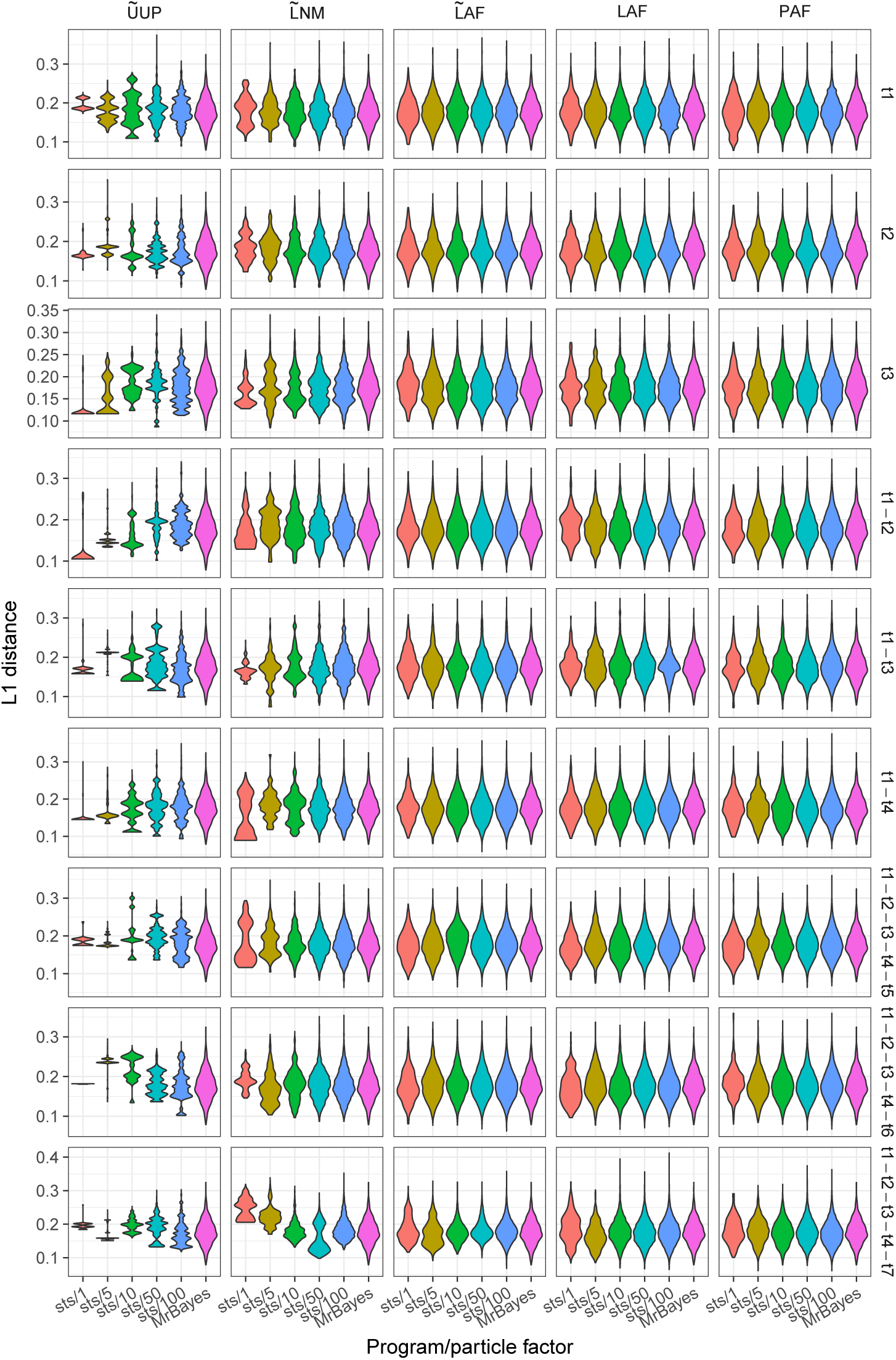
Posterior distribution of weighted Robinson-Foulds distances between each tree generated by `sts`and the maximum likelihood tree inferred by PhyML using the D2T10 data set.

**Figure S12.**
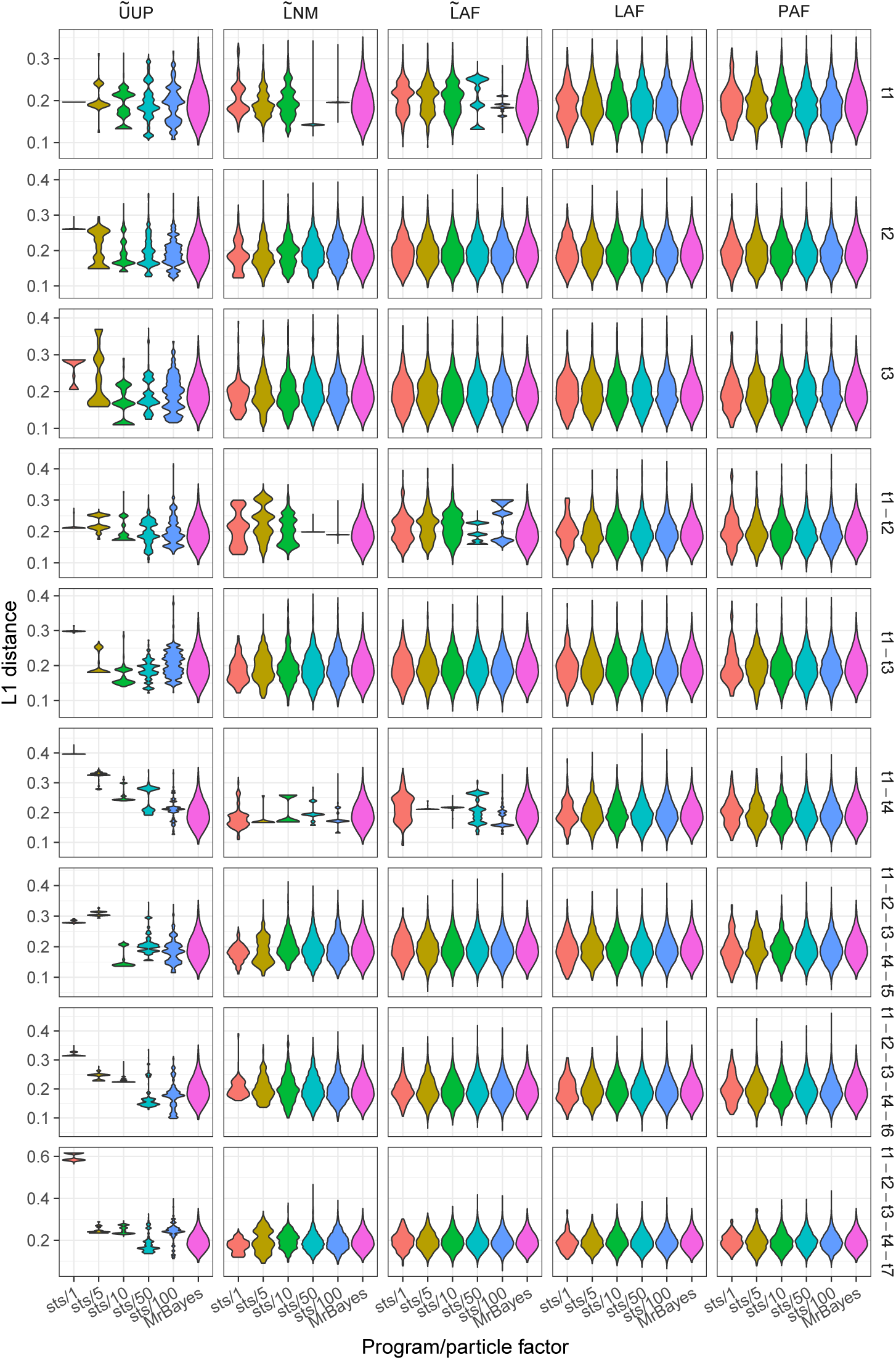
Posterior distribution of weighted Robinson-Foulds distances between each tree generated by `sts`and the maximum likelihood tree inferred by PhyML using the D3T10 data set.

**Figure S13.**
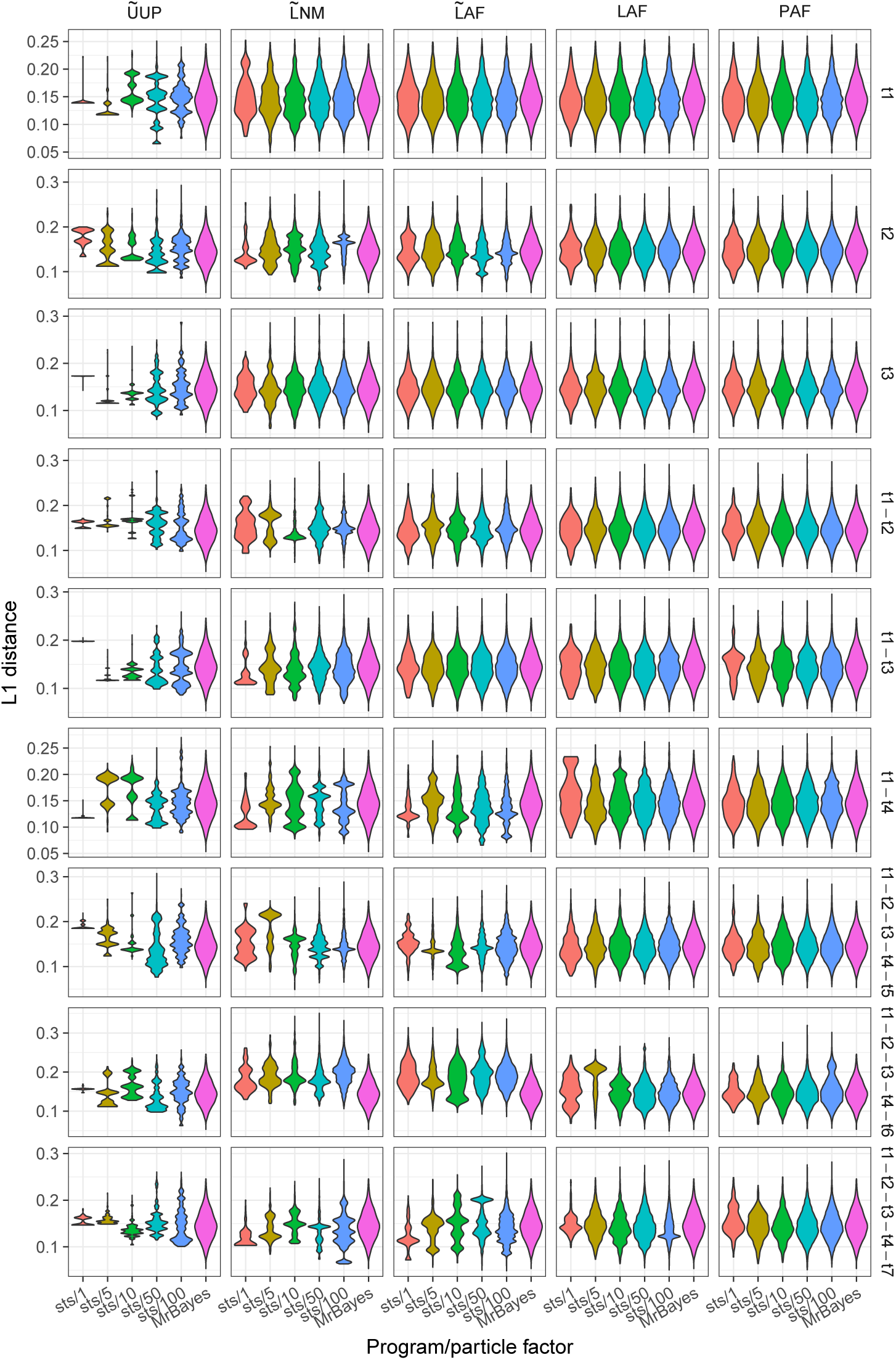
Posterior distribution of weighted Robinson-Foulds distances between each tree generated by `sts`and the maximum likelihood tree inferred by PhyML using the D4T10 data set.

**Figure S14.**
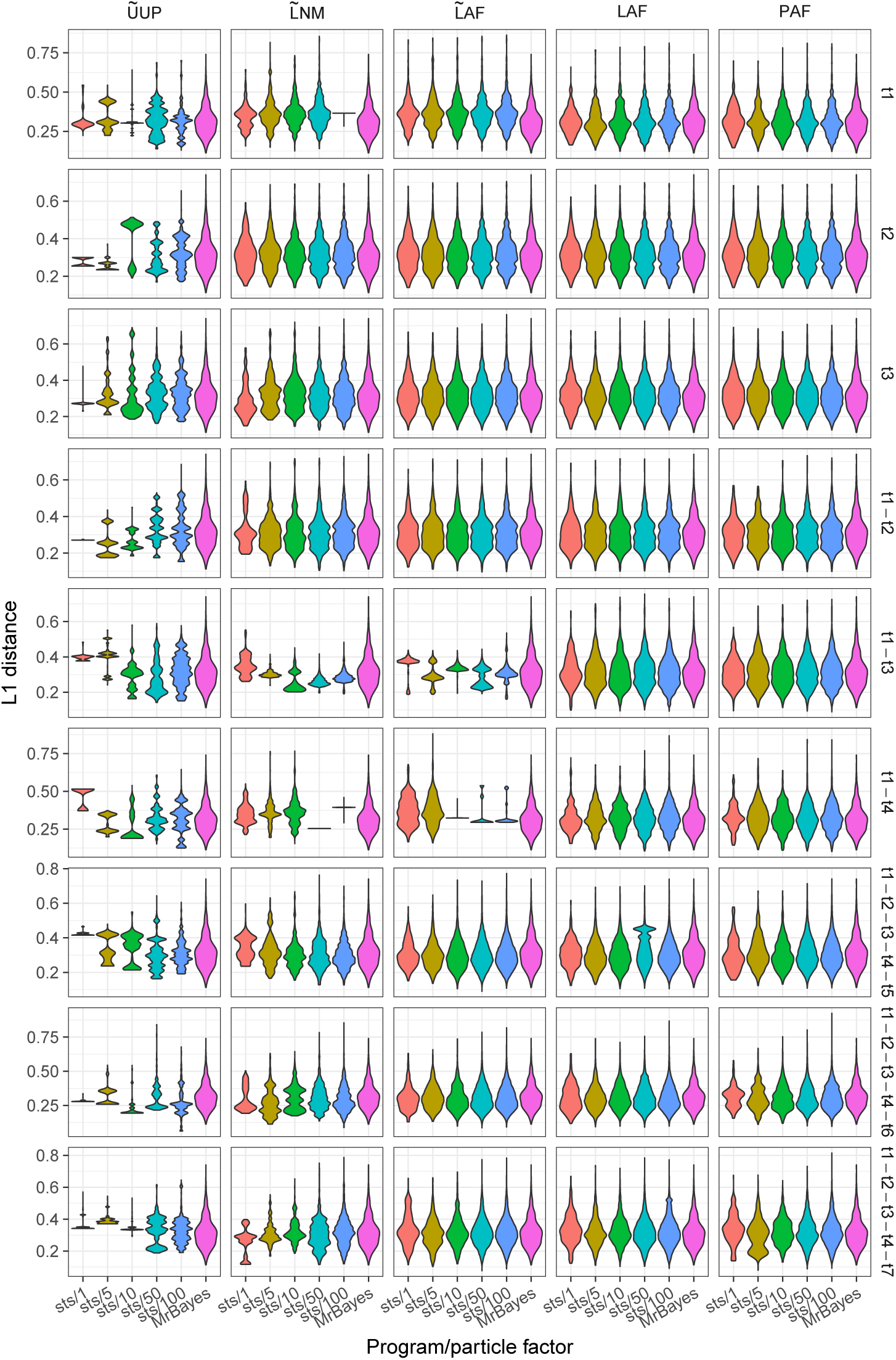
Posterior distribution of weighted Robinson-Foulds distances between each tree generated by `sts`and the maximum likelihood tree inferred by PhyML using the D5T10 data set.

**Figure S15.**
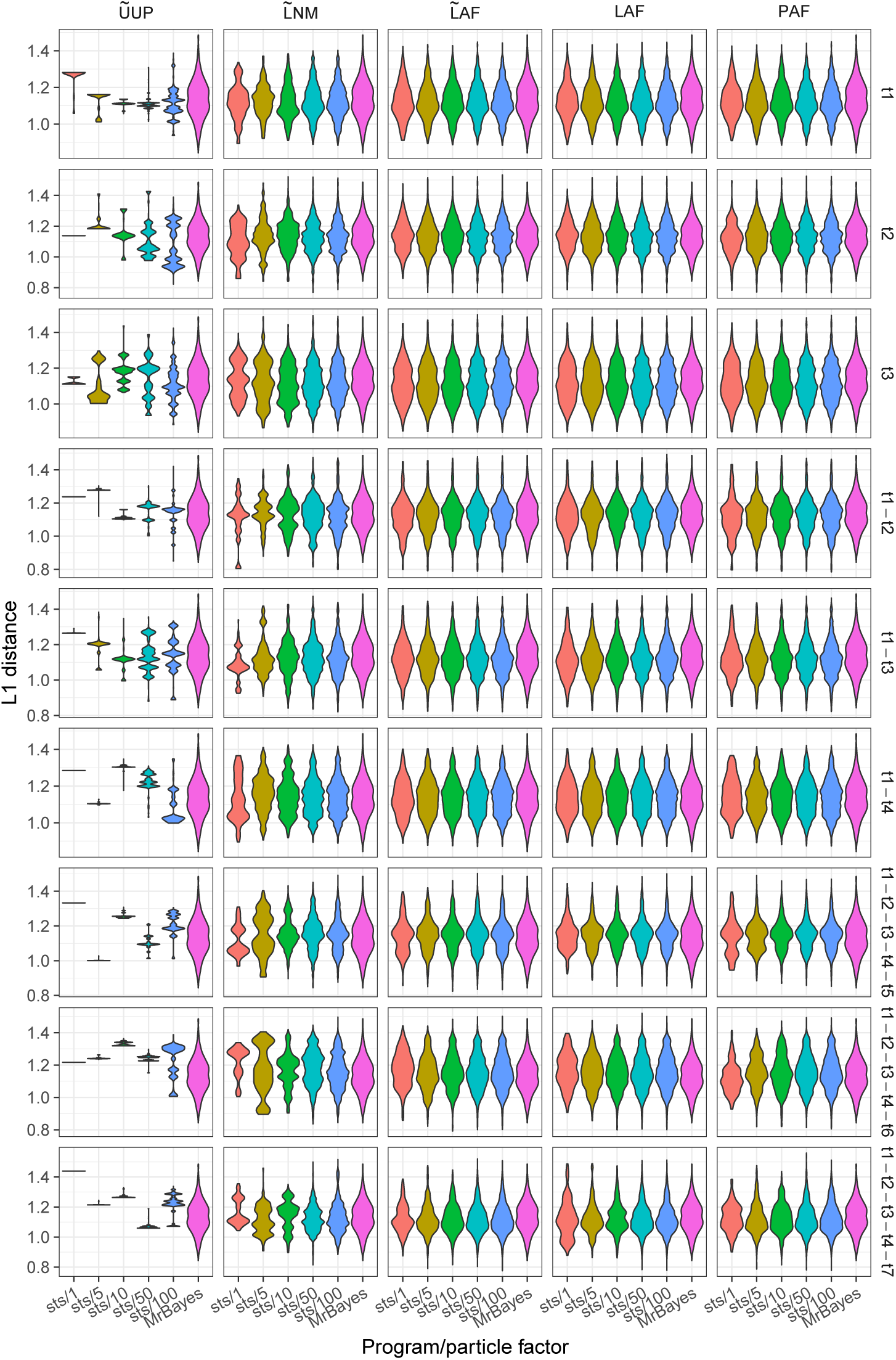
Posterior distribution of weighted Robinson-Foulds distances between each tree generated by `sts`and the maximum likelihood tree inferred by PhyML using the D1T50 data set.

**Figure S16.**
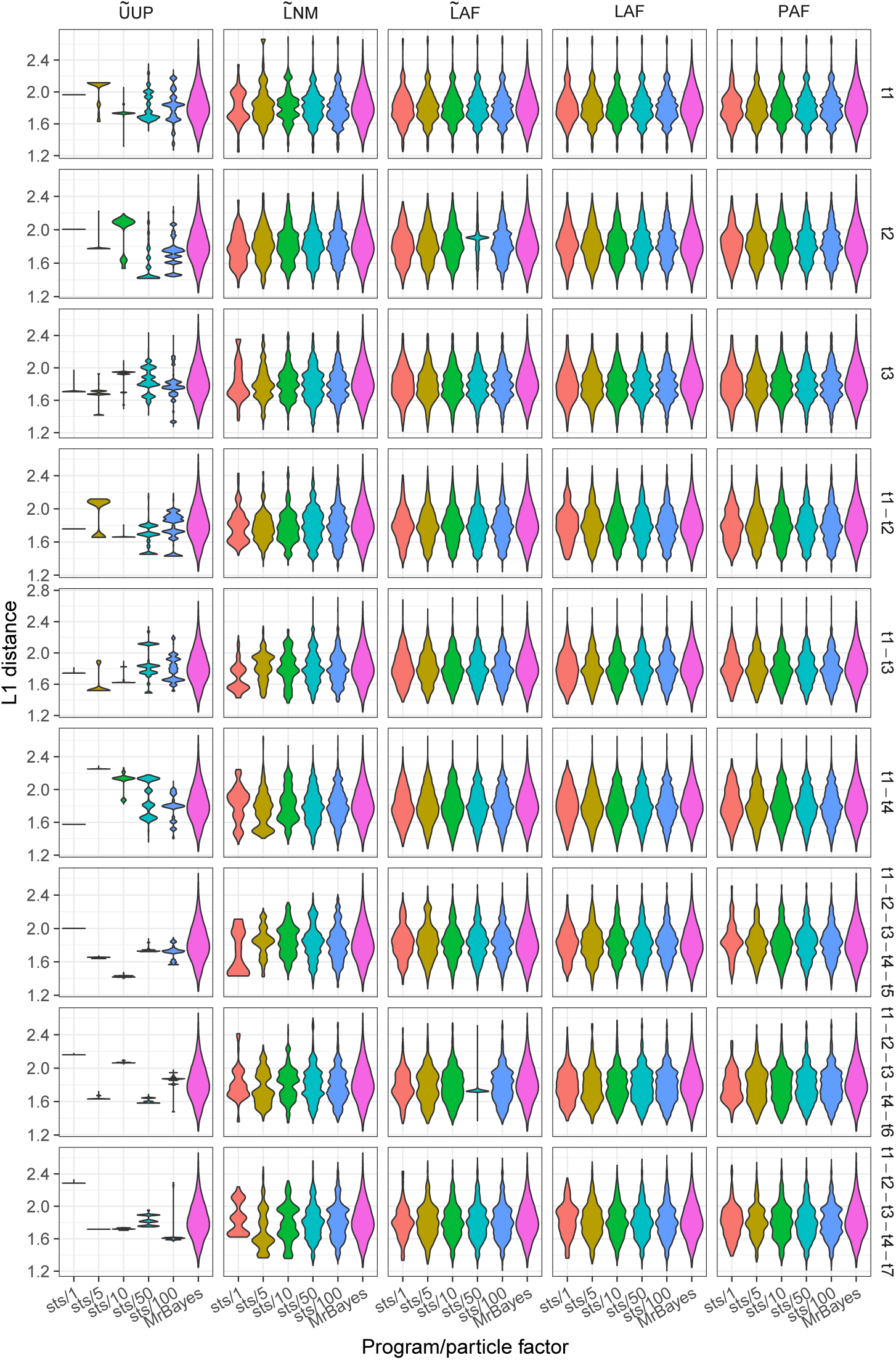
Posterior distribution of weighted Robinson-Foulds distances between each tree generated by `sts`and the maximum likelihood tree inferred by PhyML using the D2T50 data set.

**Figure S17.**
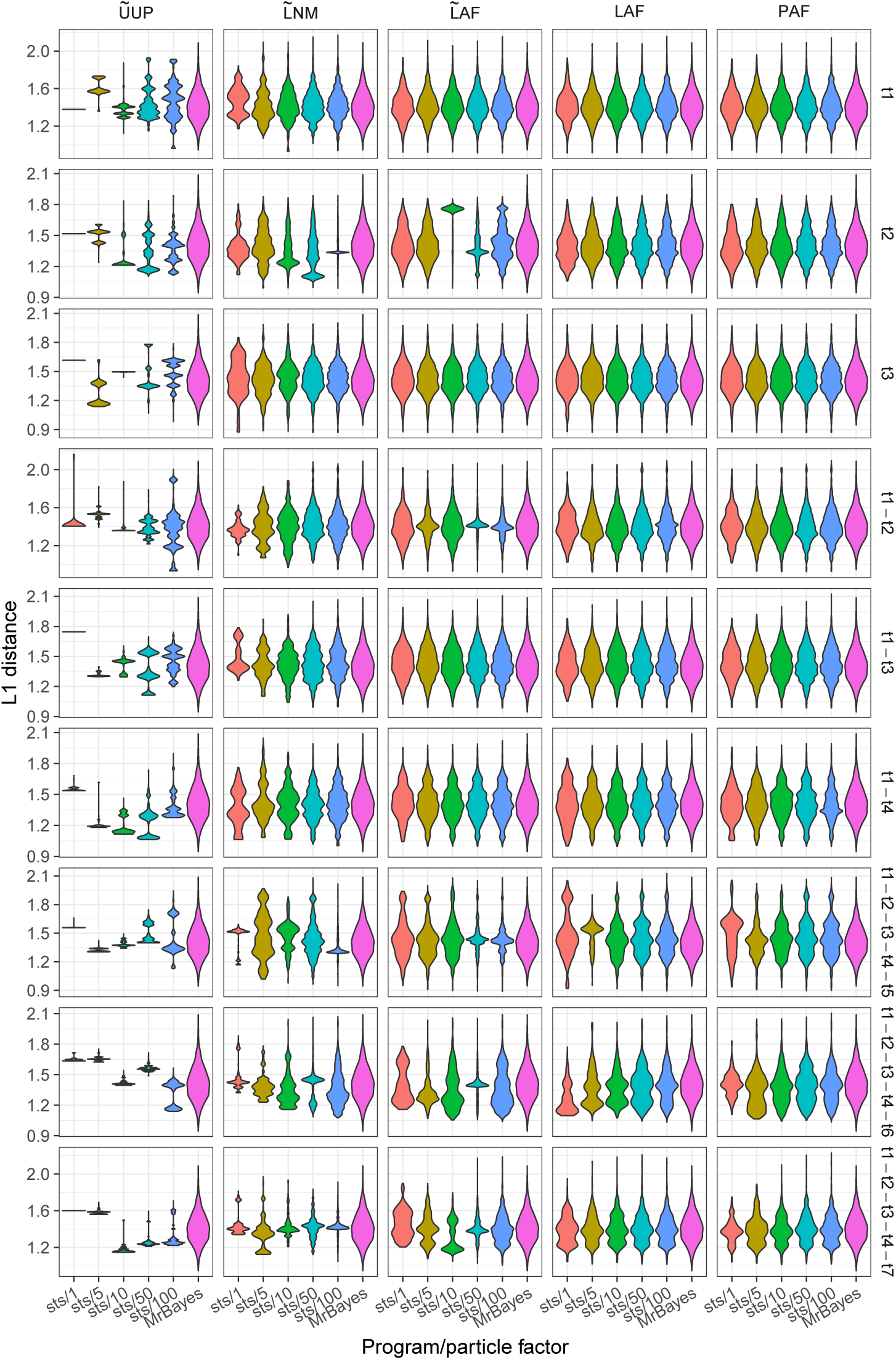
Posterior distribution of weighted Robinson-Foulds distances between each tree generated by `sts`and the maximum likelihood tree inferred by PhyML using the D3T50 data set.

**Figure S18.**
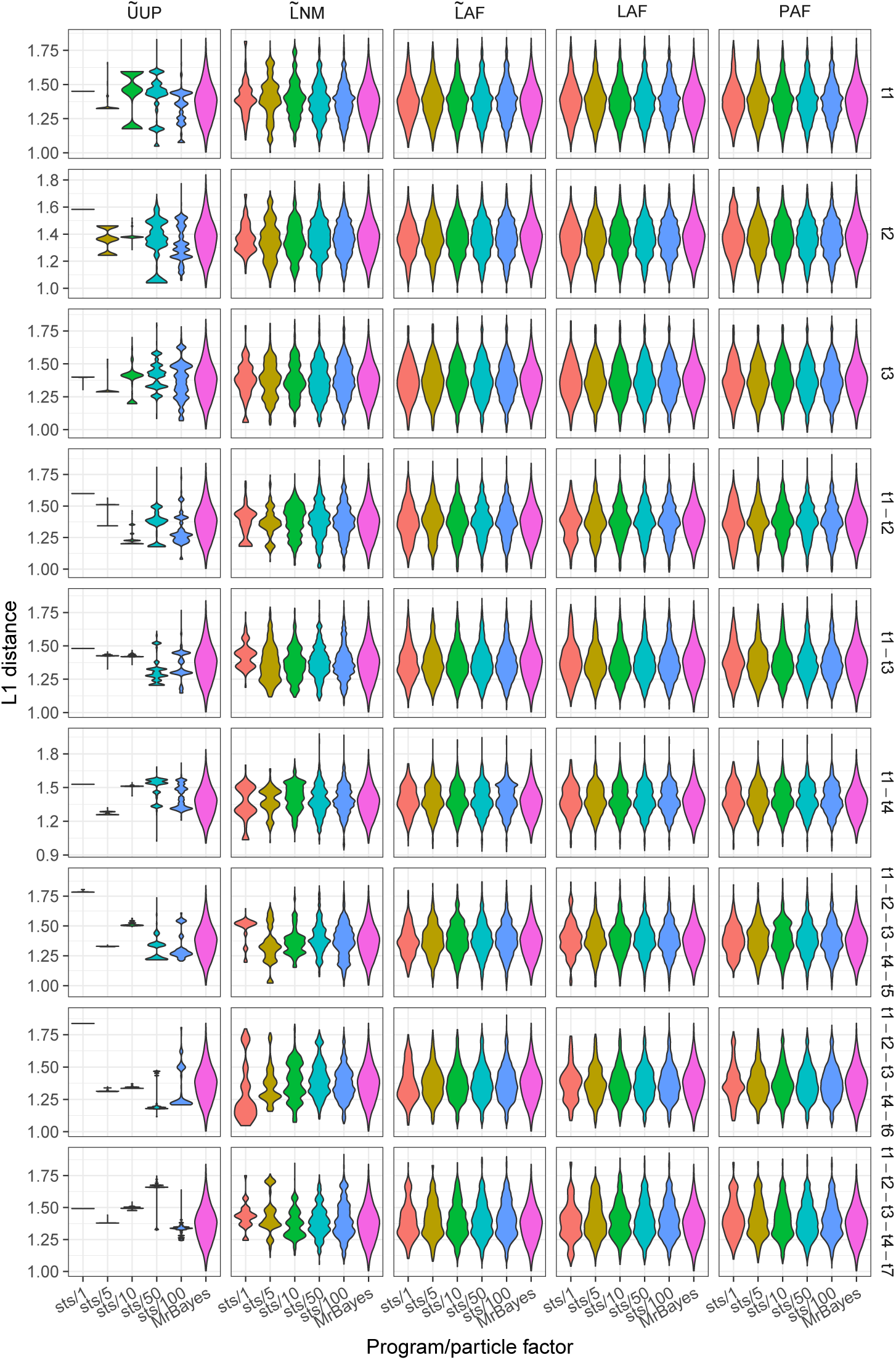
Posterior distribution of weighted Robinson-Foulds distances between each tree generated by `sts`and the maximum likelihood tree inferred by PhyML using the D4T50 data set.

**Figure S19.**
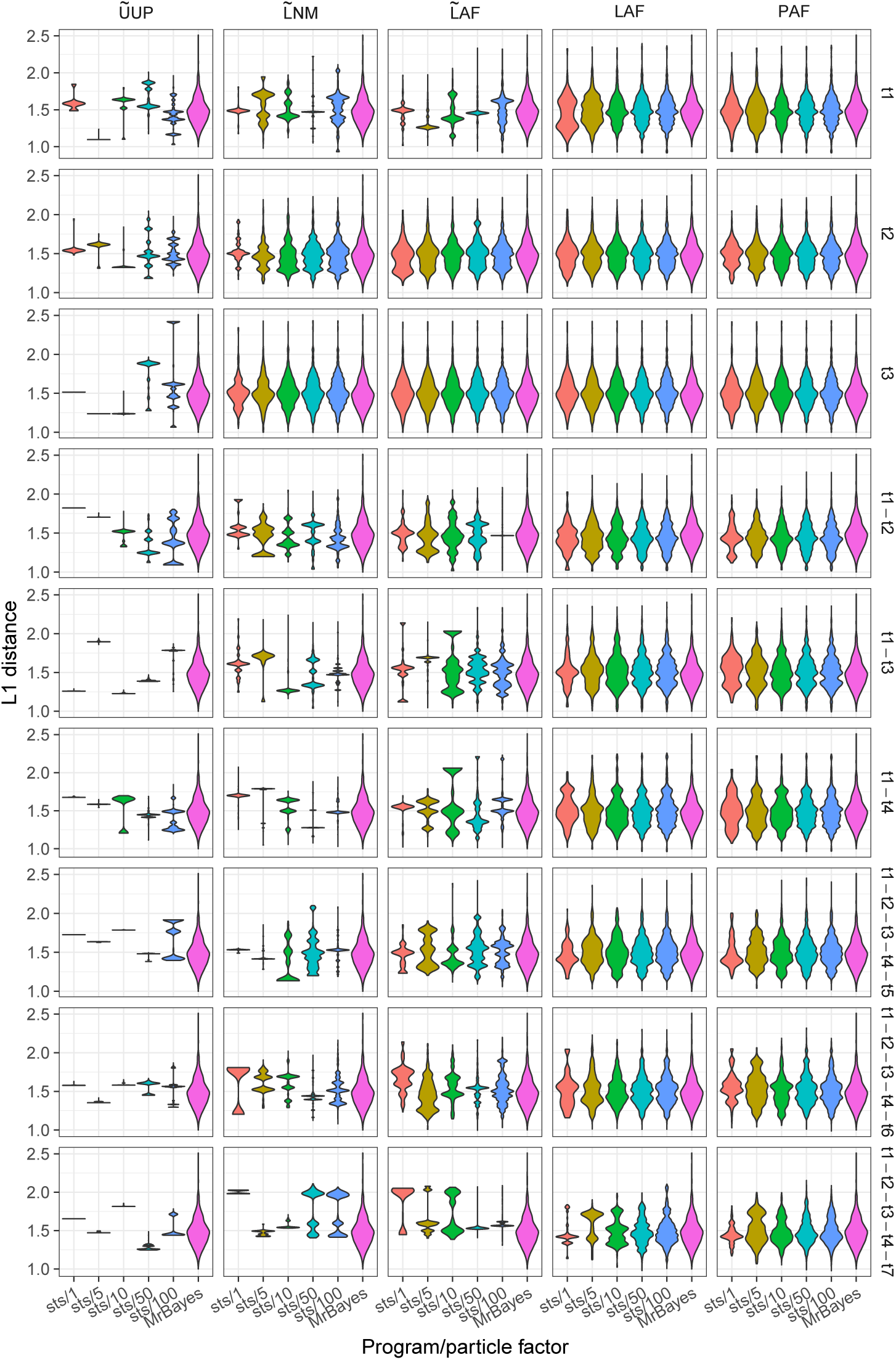
Posterior distribution of weighted Robinson-Foulds distances between each tree generated by `sts`and the maximum likelihood tree inferred by PhyML using the D5T50 data set.

**Figure S20.**
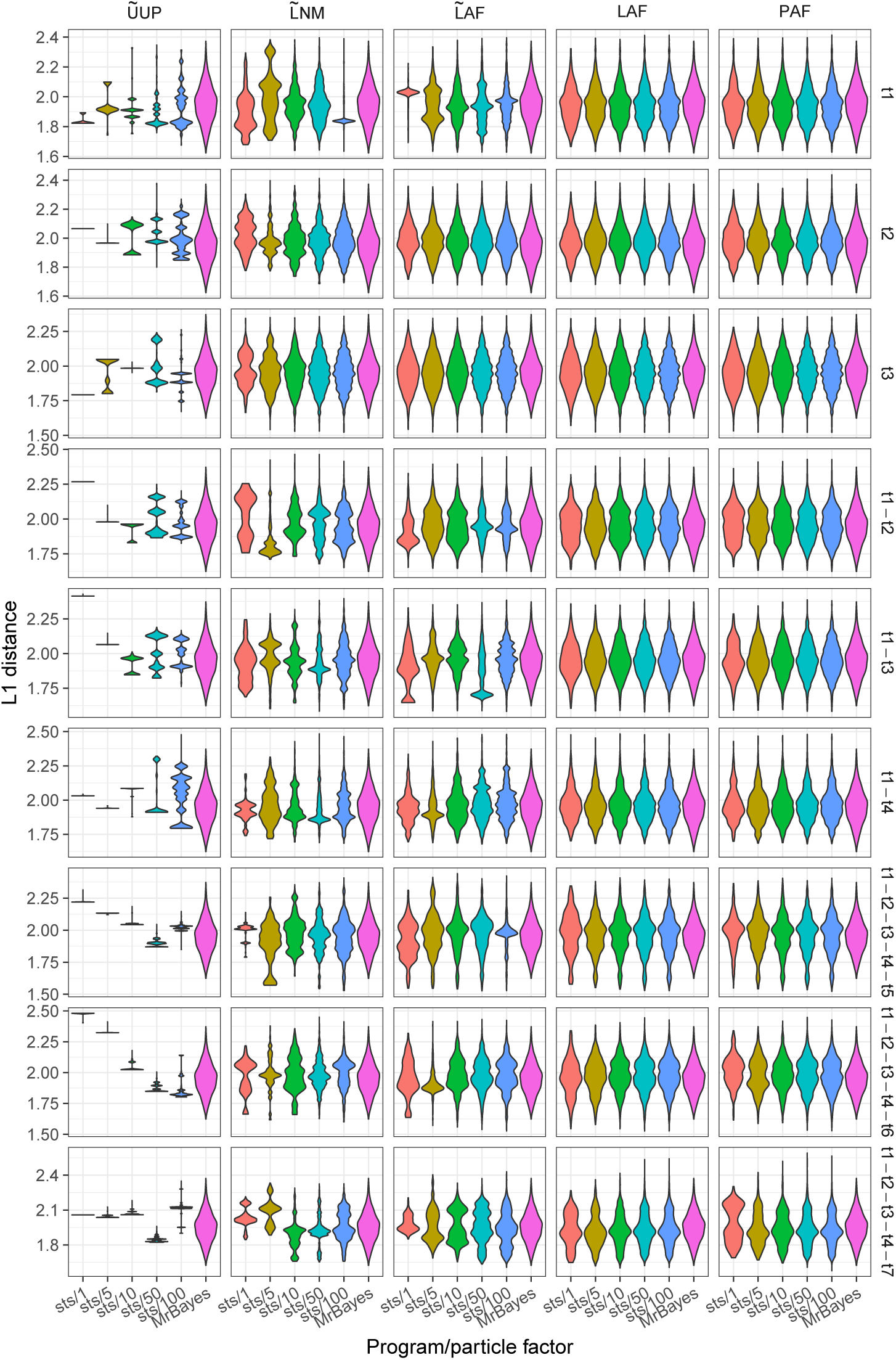
Posterior distribution of weighted Robinson-Foulds distances between each tree generated by `sts`and the maximum likelihood tree inferred by PhyML using the D1T100 data set.

**Figure S21.**
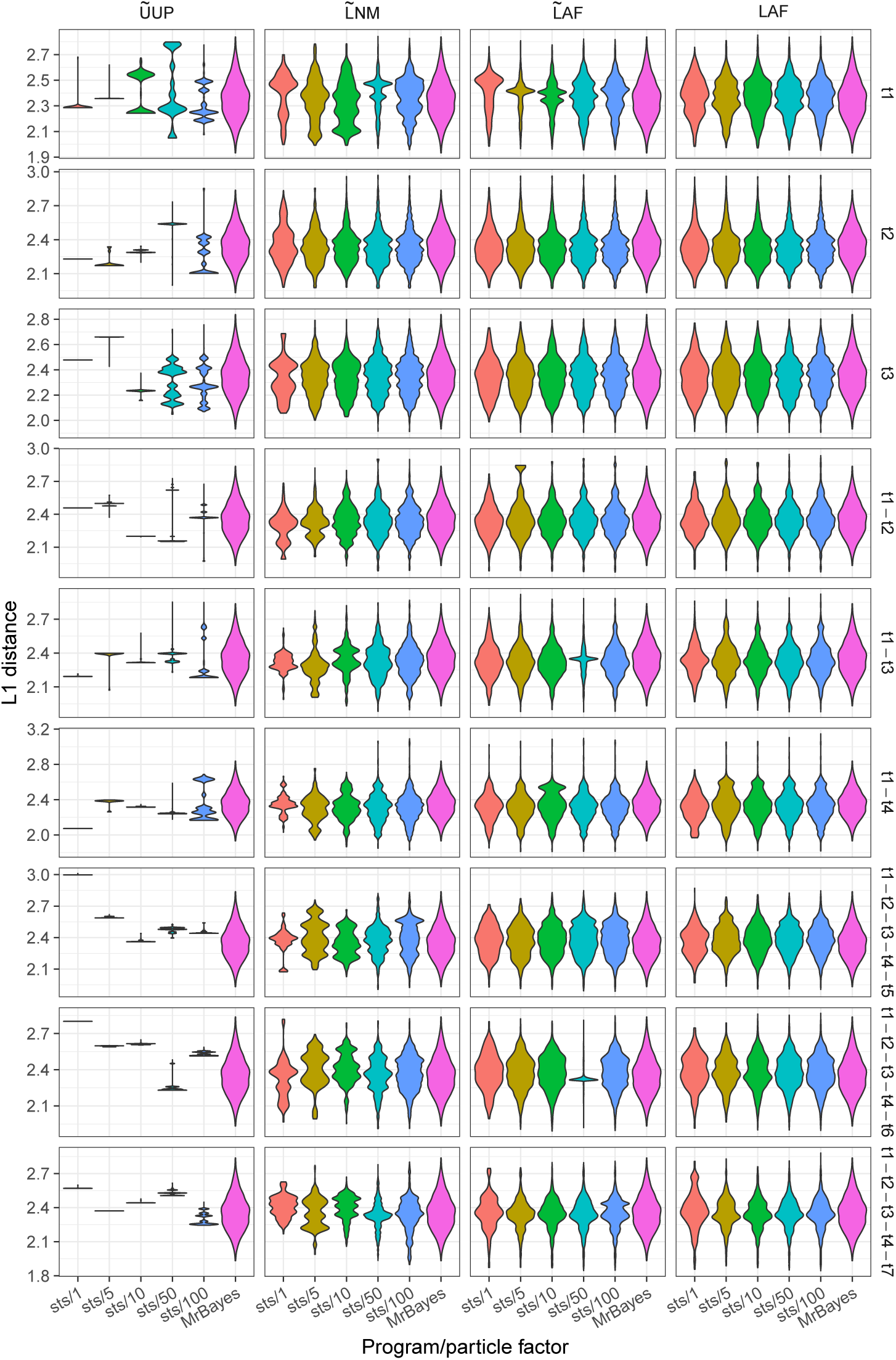
Posterior distribution of weighted Robinson-Foulds distances between each tree generated by `sts`and the maximum likelihood tree inferred by PhyML using the D2T100 data set.

**Figure S22.**
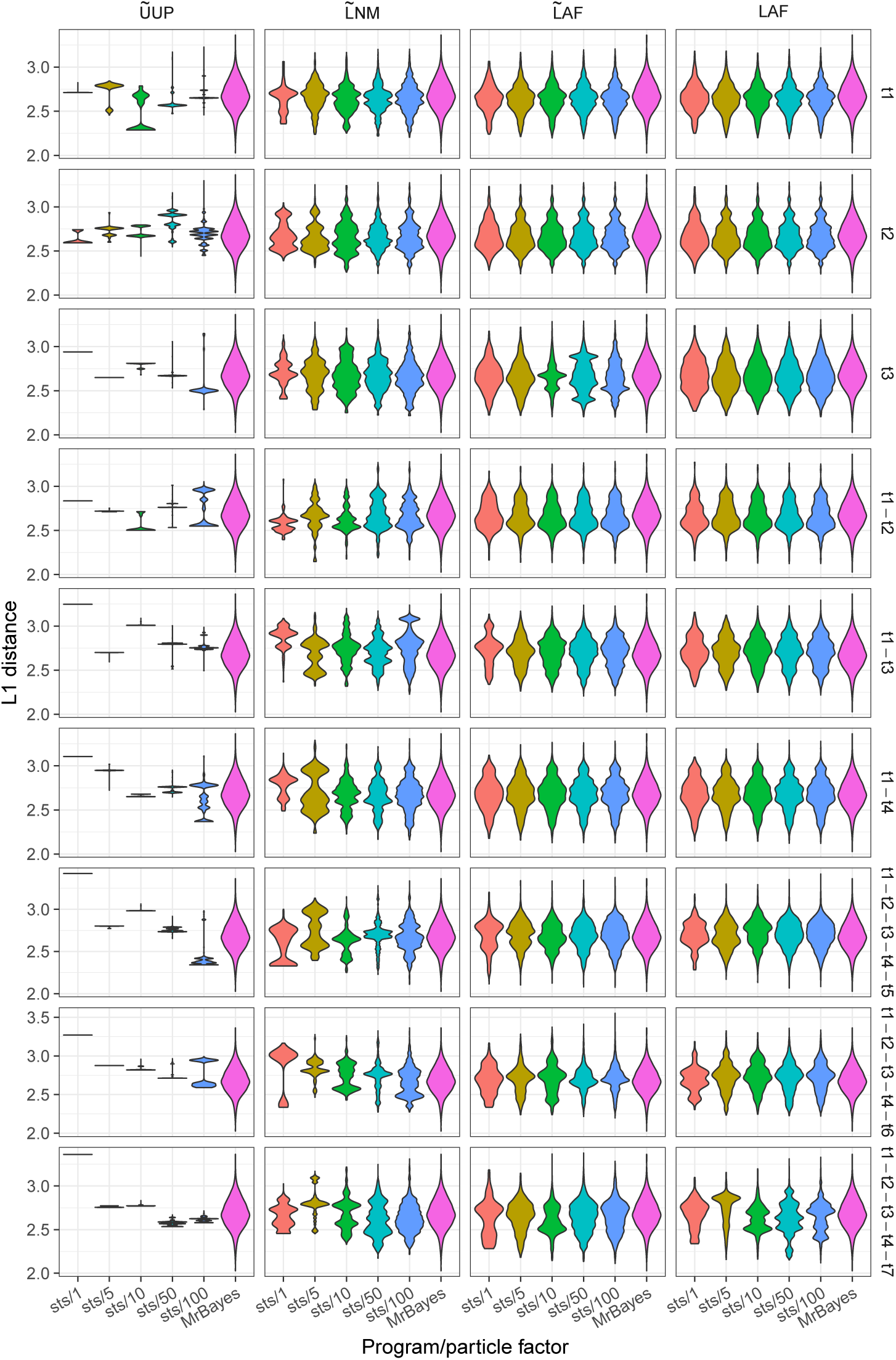
Posterior distribution of weighted Robinson-Foulds distances between each tree generated by `sts`and the maximum likelihood tree inferred by PhyML using the D3T100 data set.

**Figure S23.**
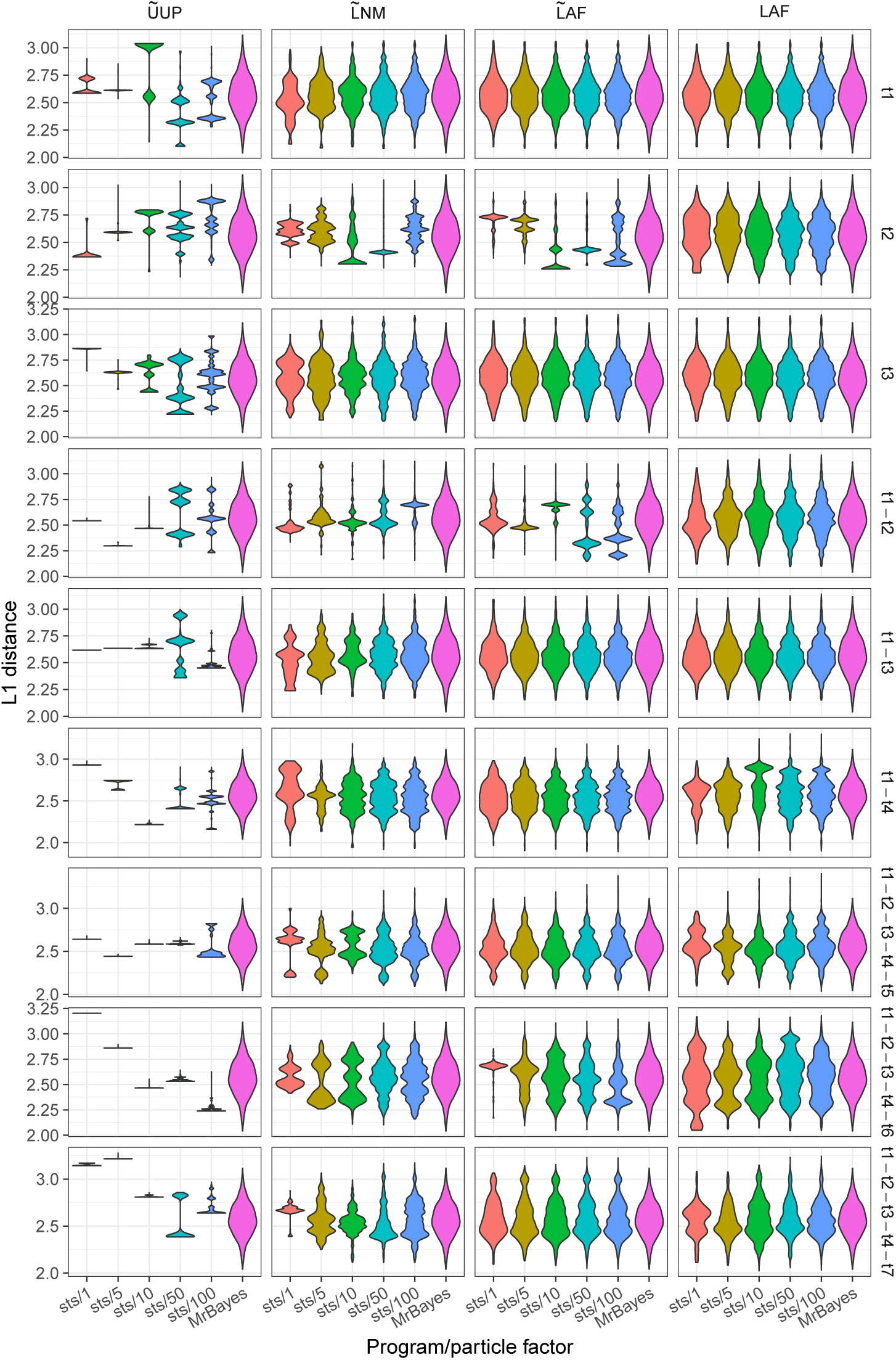
Posterior distribution of weighted Robinson-Foulds distances between each tree generated by `sts`and the maximum likelihood tree inferred by PhyML using the D4T100 data set.

**Figure S24.**
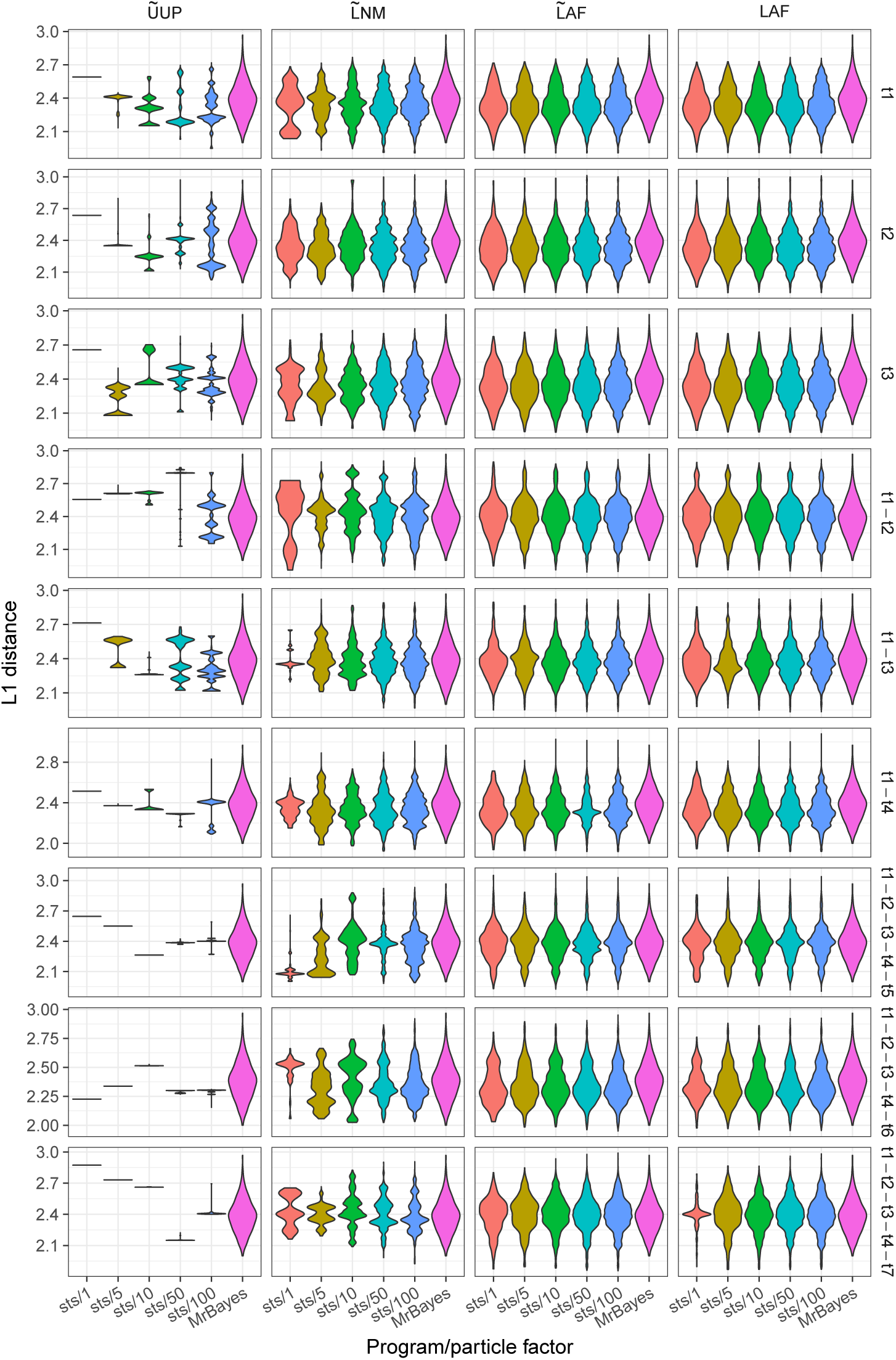
Posterior distribution of weighted Robinson-Foulds distances between each tree generated by `sts`and the maximum likelihood tree inferred by PhyML using the D5T100 data set.

